# Defining Pseudo-Haplotype Analysis Reveals Multi-Gene Genetic Pattern Across BAF Chromatin Remodeling Complexes

**DOI:** 10.64898/2026.06.22.732952

**Authors:** Xiaowei Dong, Neshatul Haque, Jessica Wagenknecht, Michael T. Zimmermann

**Affiliations:** Computational Structural Genomics Laboratory, Linda T. and John A. Mellowes Center for Genomic Sciences and Precision Medicine, Medical College of Wisconsin, Milwaukee, WI 53226, USA; Data Science Institute, Medical College of Wisconsin, Milwaukee, WI 53226, USA; Department of Biostatistics, Joseph J. Zilber College of Public Health, University of Wisconsin, Milwaukee, WI 53205, USA

**Keywords:** Pseudo-haplotypes, Chromatin remodeling, BAF (SWI/SNF) complex, Multi-variant interactions, Variant co-occurrence, Population genetics, Bioinformatics, Systems biology

## Abstract

BRG1-associated factor (BAF) is a crucial chromatin remodeling complex. Variants in genes encoding BAF complex components cause human diseases, including cancers and developmental disorders. However, the genetic diversity and variant co-occurrence patterns within BAF genes remain incompletely understood. It is feasible, though largely untested, that rare patterns of common variations could alter function similarly to rare deleterious variants. Further, there is no modern census of how often individual people simultaneously carry multiple rare and common variations, nor means for genomics practitioners to assess their combined effects. Approaches are needed to characterize complete sequences from individual samples.

In this study, we introduce a pseudo-haplotype analysis (PHA) framework, combining multiple protein-coding sequence variants, observed concurrently within individual samples, into discrete BAF patterns. In this cohort, 78.44% of pseudo-haplotype (PH) copies carry at least one BAF coding variation. Among these, 56.18% contain at least two distinct variants, and 32.39% contain three or more, indicating a substantial burden of multi-variant configurations across individuals. Notably, 25.30% of unique PHs are observed only once, highlighting a considerable proportion of people who are affected by rare or private combinations of genetic variations. We identify multiple significant (FDR < 0.05) co-occurrence combinations across global populations. These findings underscore the importance of considering population-specific genetic structures, and complete individual variant configurations when investigating disease associations and genetic mechanisms. Our approach provides a generalizable framework for characterizing multi-variant architectures within chromatin remodeling genes at a population scale, with potential applications in elucidating disease etiology and advancing precision medicine.

## Introduction

The field of genetics has made tremendous strides in understanding the human genome, yet significant challenges remain in fully interpreting the nuanced and multifaceted interplay between genetics and phenotypic expression. Recent large-scale sequencing studies further highlight the limitations of single-variant approaches. For example, rare variant analyses have shown that functionally relevant signals are highly enriched near genome-wide association studies (GWAS) loci, with enrichment reaching ∼59-fold for genes closest to GWAS sentinel variants^1^. These findings demonstrate that genetic effects are often structured, concentrate within biologically relevant regions, and motivate the development of methods capable of resolving which rare variants functionally contribute to the GWAS signal versus those that do not. Moreover, emerging evidence suggests that recombination and haplotype structure can influence the functional consequences of deleterious variants across populations^2^. Together, those observations underscore a key limitation: such analyses typically evaluate individual variants and genes rather than across the multiple gene products that assemble into protein complexes. Compellingly, these challenges not only extend to the epigenome, but they do so in part via the genes that encode epigenetic regulatory enzymes. In this way, the genome itself has the capacity to encode variant forms of chromatin regulatory enzymes that will inherently regulate the genome differently when individuals share the same exposures. Thus, we hypothesized that latent combinatorial genomic patterns may contribute to regulatory or epigenetic states that underlie human syndromes and cancer predisposition yet remain cryptic due to their presence across the multiple genes encoding a single epigenetic regulatory complex.

We term our approach, pseudo-haplotype analysis (PHA), since genetic haplotypes are sets of adjacent DNA variants that tend to be inherited together, while we seek to identify groups of germline variants that co-occur within a specific epigenetic regulatory complex. Namely, we use BAF (Bramah Associated Factor) as our vanguard example. Unlike classical haplotypes, pseudo-haplotypes (PHs) are groups of variants across continuous and non-contiguous genomic loci. In this way, we expect that genetic patterns revealed by PHA will identify cryptic relationships among genomic variations that are not apparent when analyzing variants independently.

Genetic research has simultaneously been enabled and constrained by publicly available reference sequences and gene models. The use of a common reference genome provides substantial benefits to the scientific community by establishing genomic coordinates as universal identifiers, allowing researchers to compare and integrate findings across studies. However, individuals who share genetic ancestry with the samples used to make the reference have their spectrum of normal, non-medical variation more closely matching the reference. This creates an inherent bias in genomic analysis. Recent initiatives like the Human Pangenome Reference^3^ aim to address these limitations by integrating sequences from a diverse cohort of individuals. Yet, decades of genome-wide association studies (GWAS) have successfully identified polymorphisms with strong causal or modifier effects across diverse human traits and diseases^4^. These population-level studies have revealed patterns of common genetic variation associated with complex traits. Paradoxically, clinical genetics focus on rare, highly penetrant variants while largely ignoring common variation^5,6^. This dichotomy creates a significant gap in our understanding of human genetic architecture. The ideal approach would combine both perspectives, recognizing that genetic influences on human traits and diseases exist on a continuum, and multiple variations may contribute simultaneously and non-linearly. Developing models that concurrently account for both polymorphic and individual-specific variations would enable more personalized and accurate genetic interpretation.

In this work, we develop PHA to better understand genetic variation affecting chromatin remodeling complexes and their role in human health and disease. The BAF complex provides an excellent model for studying complex genetic relationships^7^. This multi-subunit ATP-dependent chromatin remodeler plays fundamental roles in genomic regulation by altering chromatin structure around genes, BAF complexes regulate transcriptional activation and repression, influencing numerous cellular processes^7^. Several key genes encoding components of the BAF complex, including *SMARCA4* (BRG1), *SMARCA2* (BRM), *SMARCB1* (INI1), *ARID1A*, and *SMARCC1*, are essential for proper transcriptional regulation. Disruption of these genes has been implicated in a variety of human cancers, including skin, gynecologic, lower gastric, and pediatric brain tumors^8,9^, as well as congenital neurodevelopmental disorders like Coffin-Siris syndrome, autism spectrum, and more^10–12^. Understanding the complex genetics of BAF components could provide insights into shared disease mechanisms.

We posit that phenotypes result from combinations of genetic factors, including both common polymorphisms and rare variants, concurrently altering biological entities like the BAF complex, and acting in concert with environmental influences. We investigate our hypothesis using 1000 Genomes Project (1KGP)^13^ that can be accessed and verified by researchers around the globe. By examining some of the most highly studied genomes in the world, we seek an initial estimate of how common multivariate, complex-centric variant patterns are within human populations, and how discrete or individualized is the resulting landscape. We find that 94.3% of individuals carry at least one coding variant across the BAF complex, 77.5% carry ≥2 variants, and 50.9% carry ≥3 variants, with cumulative burdens ranging from 1–10 coding variants per individual. Notably, 26.5% of individuals carried ≥4 variants simultaneously, while smaller subsets exhibited substantially higher multi-variant burdens, including 11.9% with ≥5 variants and 4.97% with ≥6 variants. These findings demonstrate that combinations of common and rare variation frequently co-occur within individuals, underscoring the practical need to evaluate multi-variant configurations rather than individual variations in isolation. Additionally, there are numerous genome-wide associations already identified that can be unified according to their shared role in BAF function. Together, these findings suggest that PH approaches are needed, feasible when applied to a consistent biological entity, and will enable more accurate interpretation of individual person’s genomes.

## Material and Methods

### Genotype Data Acquisition and processing

We obtain high-coverage whole-genome sequencing Variant Call Format (VCF) files from the 1KGP Phase 3 (IGSR release 2.1), comprising 3,202 individuals sampled from across multiple continental population groups^13,14^. Genotype phasing has been performed with SHAPEIT2^15^ and post-processing using DuoHMM^16^ for chromosomes 1–22 and Eagle2 v2 for chromosome X^17^ (**Table 1**). To focus on the BAF (BRG1/BRM associated factor) chromatin remodeling complex, we retrieve genomic coordinates for 31 genes that comprise the BAF complexes, including coding exons, from Ensembl v104^18^ (**Table 2**). Using these intervals, we apply bcftools (v1.11)^19^ to subset Phase 3 VCFs. Variants are annotated with CAVA (1.2.3)^20^ to assign predicted coding functional consequences. Coding and splice-related variants with BAF complex genes, specifically missense, in-frame insertions/deletions, frame-shift, stop-gained, and splice-site variants within are retained for downstream analyses. This filtering results in 469 curated protein-coding variants across 31 BAF-complex genes.

**Table 1.**
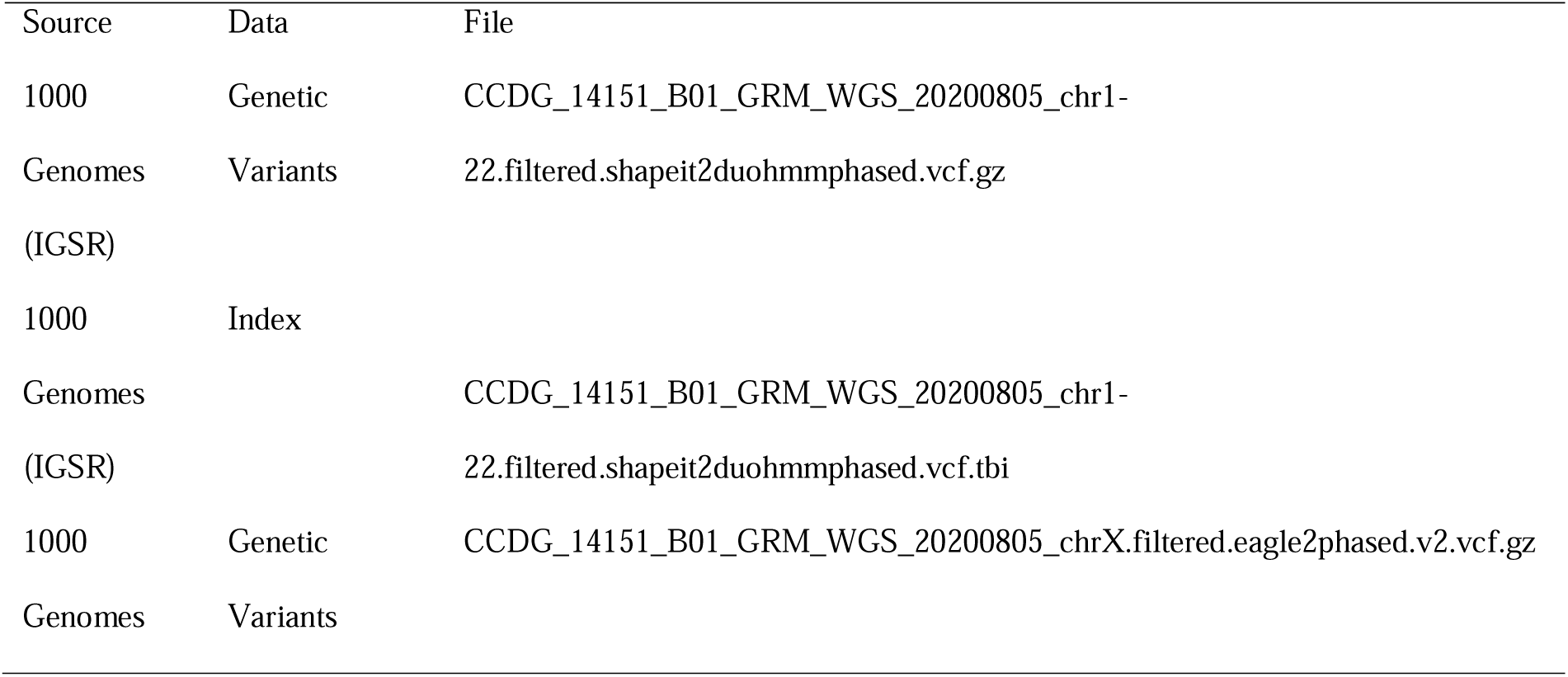
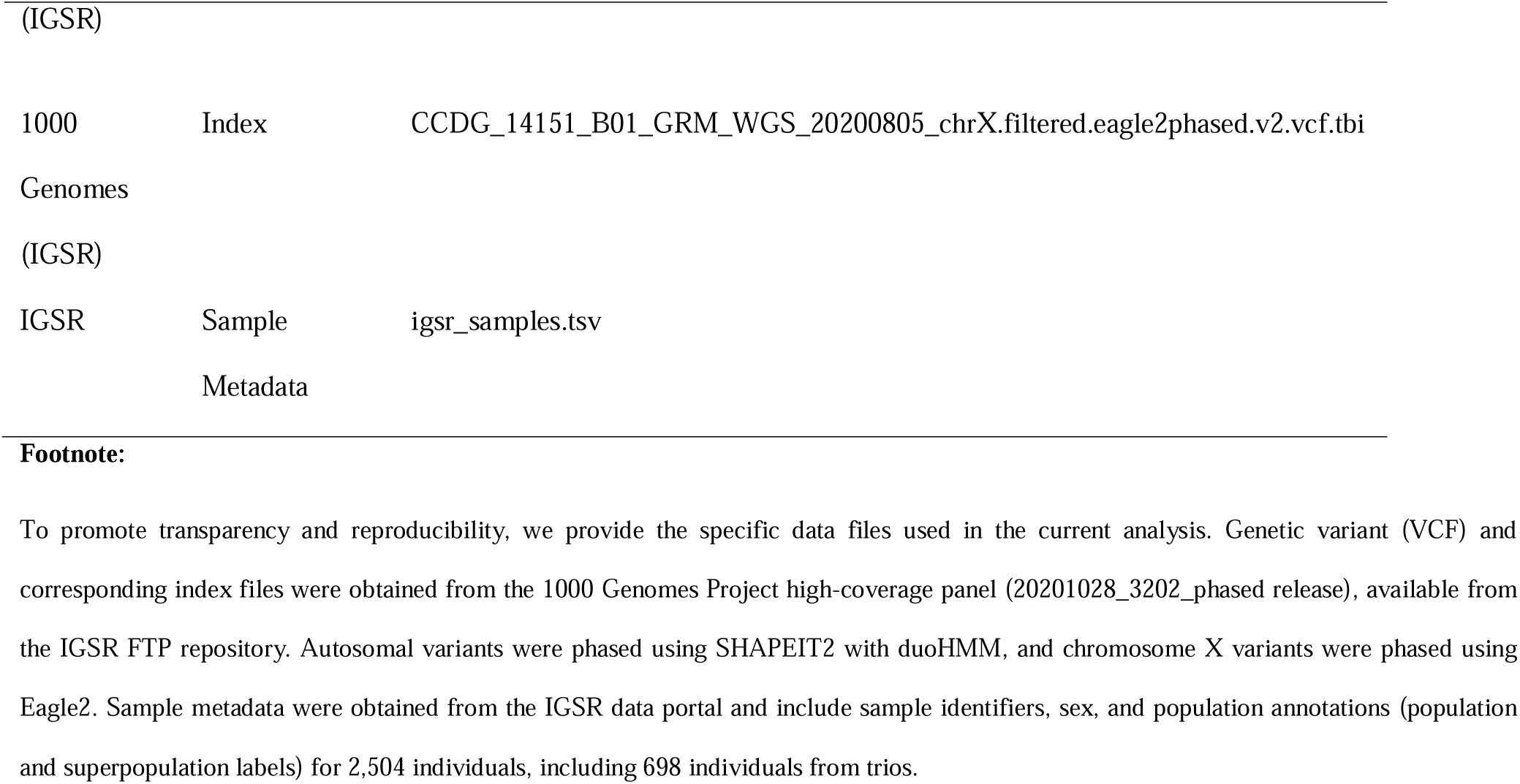
Data Source.

| Source | Data | File |
| --- | --- | --- |
| 1000 | Genetic | CCDG_14151_B01_GRM_WGS_20200805_chr1- |
| Genomes | Variants | 22.filtered.shapeit2duohmmphased.vcf.gz |
| (IGSR) |  |  |
| 1000 | Index |  |
| Genomes |  | CCDG_14151_B01_GRM_WGS_20200805_chr1- |
| (IGSR) |  | 22.filtered.shapeit2duohmmphased.vcf.tbi |
| 1000 | Genetic | CCDG_14151_B01_GRM_WGS_20200805_chrX.filtered.eagle2phased.v2.vcf.gz |
| Genomes | Variants |  |
| (IGSR) |  |  |
| 1000 | Index | CCDG_14151_B01_GRM_WGS_20200805_chrX.filtered.eagle2phased.v2.vcf.tbi |
| Genomes |  |  |
| (IGSR) |  |  |
| IGSR | Sample | igsr_samples.tsv |
|  | Metadata |  |

To promote transparency and reproducibility, we provide the specific data files used in the current analysis. Genetic variant (VCF) and corresponding index files were obtained from the 1000 Genomes Project high-coverage panel (20201028\_3202\_phased release), available from the IGSR FTP repository. Autosomal variants were phased using SHAPEIT2 with duoHMM, and chromosome X variants were phased using Eagle2. Sample metadata were obtained from the IGSR data portal and include sample identifiers, sex, and population annotations (population and superpopulation labels) for 2,504 individuals, including 698 individuals from trios.

**Table 2.**
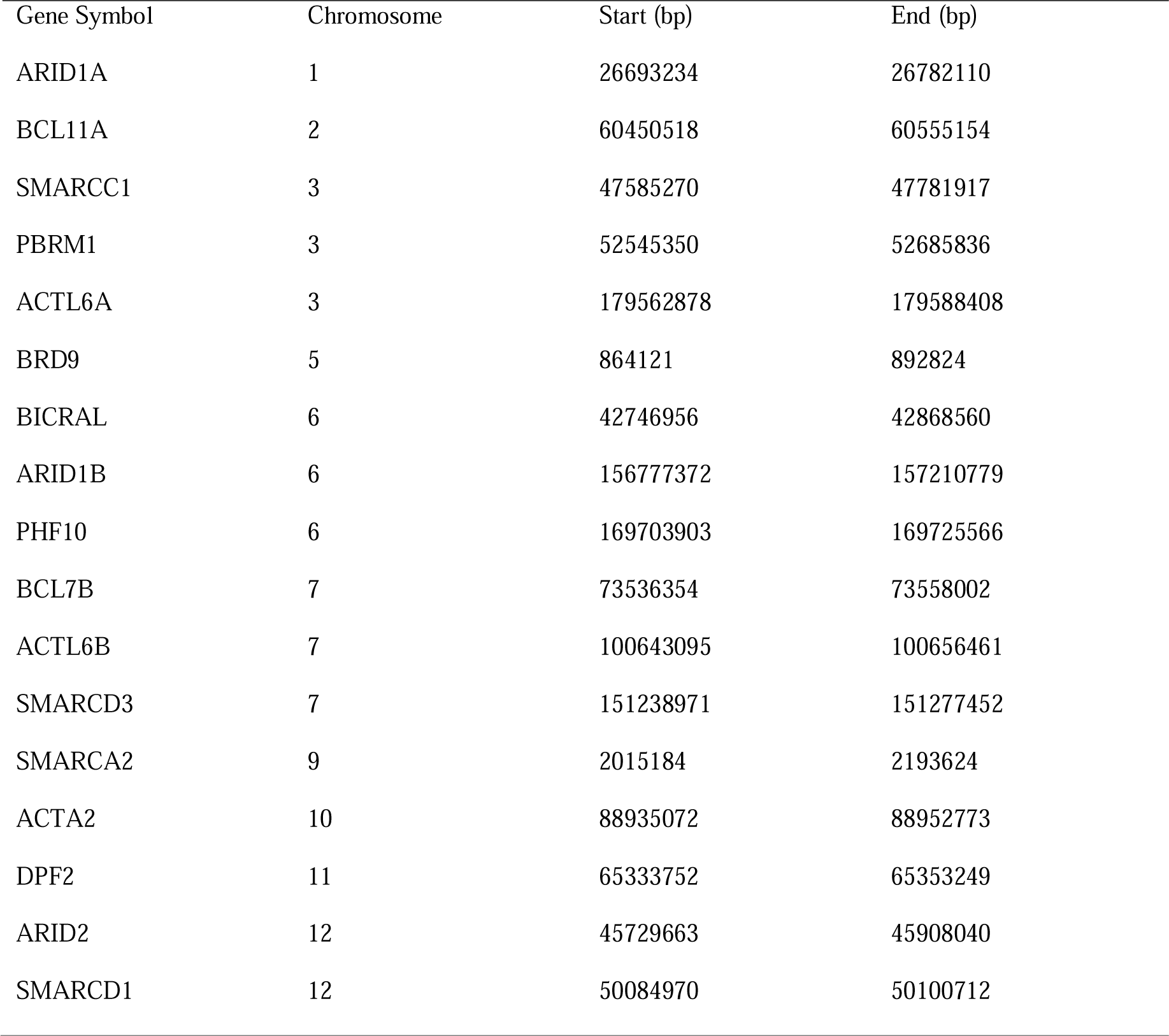
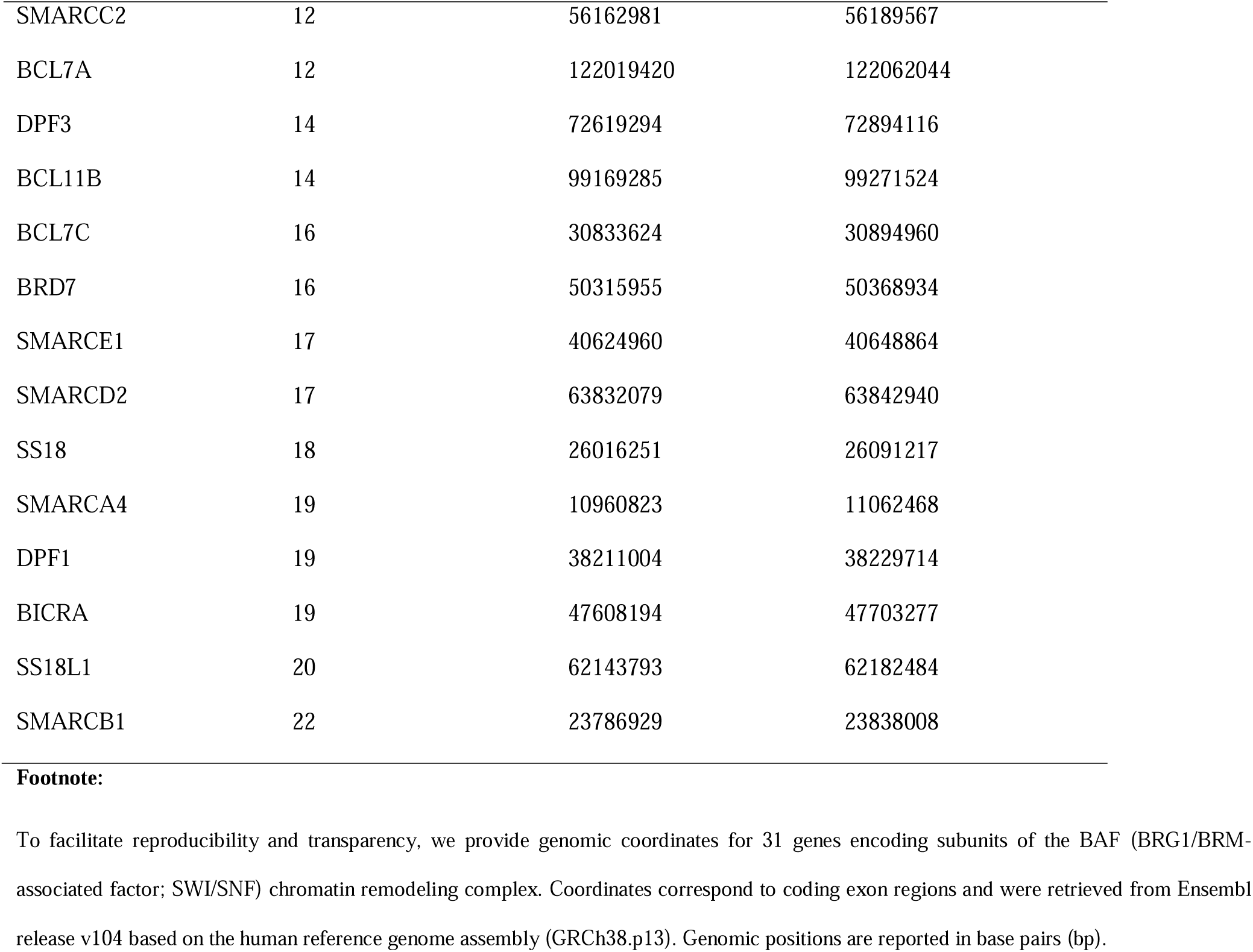
Genomic coordinates of BAF chromatin remodeling complex genes.

| Gene Symbol | Chromosome | Start (bp) | End (bp) |
| --- | --- | --- | --- |
| ARID1A | 1 | 26693234 | 26782110 |
| BCL11A | 2 | 60450518 | 60555154 |
| SMARCC1 | 3 | 47585270 | 47781917 |
| PBRM1 | 3 | 52545350 | 52685836 |
| ACTL6A | 3 | 179562878 | 179588408 |
| BRD9 | 5 | 864121 | 892824 |
| BICRAL | 6 | 42746956 | 42868560 |
| ARID1B | 6 | 156777372 | 157210779 |
| PHF10 | 6 | 169703903 | 169725566 |
| BCL7B | 7 | 73536354 | 73558002 |
| ACTL6B | 7 | 100643095 | 100656461 |
| SMARCD3 | 7 | 151238971 | 151277452 |
| SMARCA2 | 9 | 2015184 | 2193624 |
| ACTA2 | 10 | 88935072 | 88952773 |
| DPF2 | 11 | 65333752 | 65353249 |
| ARID2 | 12 | 45729663 | 45908040 |
| SMARCD1 | 12 | 50084970 | 50100712 |
| SMARCC2 | 12 | 56162981 | 56189567 |
| BCL7A | 12 | 122019420 | 122062044 |
| DPF3 | 14 | 72619294 | 72894116 |
| BCL11B | 14 | 99169285 | 99271524 |
| BCL7C | 16 | 30833624 | 30894960 |
| BRD7 | 16 | 50315955 | 50368934 |
| SMARCE1 | 17 | 40624960 | 40648864 |
| SMARCD2 | 17 | 63832079 | 63842940 |
| SS18 | 18 | 26016251 | 26091217 |
| SMARCA4 | 19 | 10960823 | 11062468 |
| DPF1 | 19 | 38211004 | 38229714 |
| BICRA | 19 | 47608194 | 47703277 |
| SS18L1 | 20 | 62143793 | 62182484 |
| SMARCB1 | 22 | 23786929 | 23838008 |

To facilitate reproducibility and transparency, we provide genomic coordinates for 31 genes encoding subunits of the BAF (BRG1/BRM-associated factor; SWI/SNF) chromatin remodeling complex. Coordinates correspond to coding exon regions and were retrieved from Ensembl release v104 based on the human reference genome assembly (GRCh38.p13). Genomic positions are reported in base pairs (bp).

### Cohort Metadata Processing and Super-Population Harmonization

Sample metadata including sex and population labels are obtained from the sample information files of the 1KGP Phase 3 hosted by the International Genome Sample Resource (IGSR)^13,14^.To ensure consistency and adequate sample sizes for statistical analysis, population labels are harmonized into five continental-level super-populations: African (AFR), American (AMR), East Asian (EAS), European (EUR), or South Asian (SAS). Ancestry assignments from the 1KGP are used as the primary framework, and reference-based population labels derived from the Simons Genome Diversity Project (SGDP)^21^ are subsequently mapped to their corresponding 1KGP continental super-populations as follows: African Ancestry and Africa samples are grouped into AFR super-population; East Asian Ancestry and East Asia into EAS; South Asian Ancestry and South Asia into SAS; and European Ancestry samples are combined with European-associated admixed categories into EUR. Samples labeled as American Ancestry are retained as a distinct AMR super-population. The resulting harmonized super-population labels are merged with the filtered genotype dataset to enable population-stratified and ancestry-aware downstream analyses.

### GWAS Integration and SNP–Gene–Trait Mapping

Genome-wide association study (GWAS) data are obtained from the NHGRI-EBI GWAS^4^ Catalog to assess population-level disease associations across 31 BAF-complex genes. A total of 1,343 unique GWAS risk alleles associated with 904 traits are compiled and integrated with 469 curated protein-coding variants derived from the 1KGP. To prioritize functionally relevant associations, GWAS SNPs are filtered based on proximity to protein-coding variants within a ±10 kb window, approximating local linkage disequilibrium (LD) structure^22^. SNPs are mapped to nearby coding variants to generate SNP–gene–trait associations. Genes lacking overlapping GWAS signals (*ACTA2* and *ACTL6A*) are excluded, resulting in a refined set of 29 genes. Shared traits are defined as traits associated with SNPs mapping two or more BAF-complex genes.

### Allele Frequency Comparison Between 1KGP and gnomAD

To evaluate the robustness of allele frequency estimates, variant allele frequencies derived from the 1KGP are compared with those reported in gnomAD v4.1.0^23^. Variants are matched between datasets based on genomic position and allelic identity. Only variants successfully matched across both datasets are retained for comparison. Correlation analysis was performed on log_10_-transformed allele frequencies.

### Pseudo-Haplotype Construction

To investigate combinatorial patterns of genetic variation across the BAF complex, we construct PHs by decomposing phased diploid genotypes from 3,202 individuals in the 1KGP. Each genotype is represented as a phased biallelic sequence and split into its two constituent haploid genomes. This process yields 6,404 chromosomal PH copies for downstream analysis. PHs are defined by aggregating the carrier status (presence/absence) of curated protein-altering variants across 31 BAF-complex genes (469 loci). Each PH copy represents a haploid genome with a specific variant configuration. PH copies sharing identical variant configurations are collapsed into distinct PHs. The PH recurrence level is defined as the number of PH copies observed for each distinct PH. The PH frequency of each PH is calculated as its proportion among all 6,404 PH copies.

Because parental origin information is unavailable, PH copies could not be assigned to consistent maternal or parental chromosomes. Therefore, analyses are performed at both PH copy level and individual level resolutions. For individual level analyses, each protein-coding variant is considered present if detected in either PH copies for a given individual.

### PH Frequency-based Filtering

To distinguish recurrent population-level patterns from rare or private variation, we apply frequency-based filtering to PHs. PHs observed at a frequency below 0.05% PH frequency (corresponding to <4 of 6,404 PH copies; recurrence level <4) are classified as rare, whereas those observed at ≥0.05% PH frequency (≥4 of 6,404 PH copies; recurrence level ≥4) are retained as common for analyses of recurrent PH configurations. This empirical PH frequency threshold is selected to reduce the influence of singleton and ultra-low-recurrence haplotypes, which are more likely to reflect private, population-specific, or potentially unstable configurations, while retaining reproducibly observed PHs across samples.

### Variant Burden and Recurrence in BAF-Complex Pseudo-Haplotypes

To characterize the distribution and relationship between recurrence and combinatorial complexity of PHs, we analyze protein-coding variant configurations across all PH copies. Variant burden per PH is defined as the number of protein-coding variants present within each PH configuration. Distributions of PH recurrence levels and variant burden are summarized across all PH copies to assess how variant complexity relates to PH frequency. We generate PH similarity trees and Tanghulu-style presence versus absence visualizations based on protein-coding variant configurations. Pairwise similarity between PHs is quantified using the Jaccard index for construction of the similarity tree, and variant co-occurrence patterns are visualized using Tanghulu-style plots. Analyses are restricted to PHs meeting the predefined PH frequency threshold.

### Structural Mapping of Protein-Coding Variants

Protein-coding variants are mapped onto a three-dimensional model of the canonical BAF (cBAF) complex comprising 10 core subunits (SMARCA4, ARID1A, SMARCC1, SMARCC2, SMARCD1, SMARCE1, ACTB, ACTL6A, SMARCB1, and DPF2), along with associated DNA and histone components. A subset of variants (n = 118) is selected from the total set of 469 curated variants that map to structurally resolved regions in this specific sub-complex configuration. Variants within unresolved regions (n = 81) are mapped to their N-terminal-most resolved amino acid while those resolved (n = 37) are directly mapped to atomic coordinates. Structural visualization was performed using PyMOL v3.1.8^24^.

### Statistical Analysis of Multi-variant Enrichment

To quantify combinatorial interactions among protein-coding variants within BAF complex genes, we perform systematic co-occurrence and mutual exclusivity analyses using phased genotype data from 3,202 individuals in the 1KGP^13,14^. Phased diploid genotypes are decomposed into PH copies for downstream analyses. Each PH copy is encoded as a binary vector across variant loci, where:

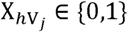

denotes the allelic state of variant V*_j_* on PH copy *h* (0 = reference allele, 1 = alternative allele). A binary genotype matrix is constructed with PH copies as rows and variant loci as columns. Analyses are restricted to variants present meeting the predefined PH frequency threshold (≥0.05%) described in *Pseudo-Haplotype Construction* and *Frequency-based Filtering* to reduce instability from sparse counts. For multi-variant enrichment analyses, only PH copies carrying two or more alternative alleles within the evaluated set are subsequently retained. Marginal allele frequencies and independence expectations are calculated using all 6,404 PH copies before this filtering step. For each combination of *k* variants (*k* = 2, 3,…, 10), including both pairwise and higher-order combinations, we quantify observed co-occurrence, marginal allele frequencies, expected co-occurrence under an independence model, and deviation from independence using a log_2_ enrichment (LEC) statistic.

#### Observed Co-Occurrence

For a set of *k* variants V*_1_*, V*_2,_*……V*_k_*, the observed number of PH copies carrying all *k* alternative alleles is defined as:

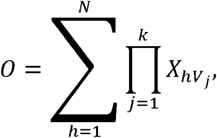

where N is the total number of PH copies.

#### Marginal Allele Frequencies

The marginal alternative-allele frequency for variant V*_j_* is computed as:

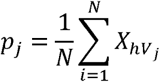

#### Expected Co-Occurrences Under Independence

Assuming independence among variants, the expected number of PH copies carrying all *k* alternative alleles is calculated as:

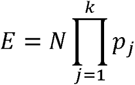

#### Log_2_ Enrichment (LEC) of Co-Occurrences

Deviation from independence is quantified using log_2_ Enrichment (LEC) statistic comparing observed to expected co-occurrences counts. The LEC statistic is analogous to a log_2_-transformed odds ratio and quantifies directional deviation from independence while preserving symmetry between enrichment and depletion:

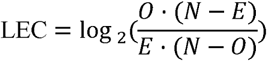

To ensure numerical stability, a pseudo-count of 0.5 is added to *O*, *N* – *O*, *E*, and *N* – *E*, when necessary.

#### Statistical Significance Testing

For each variant combination with *k* ≥2, statistical significance of deviation from independence is evaluated using a two-sided exact binomial test under the null model:

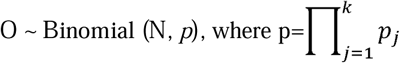

Resulting p-values are corrected for multiple hypothesis testing using the Benjamini–Hochberg false discovery rate (FDR) procedure^25^.

#### Interpretation of Multi-variant Enrichment Effect

Variant combinations are classified as co-occurring when FDR < 0.05 and LEC > 0, mutually exclusive when FDR< 0.05 and LEC < 0, and not significant when FDR ≥ 0.05. This haplotype-based framework enable detection of multi-variant configurations that are statistically enriched or depleted relative to expectations under marginal allele independence.

### Global and Population-Stratified Co-Occurrence Patterns of Multi-Variant Pseudo-Haplotypes

To evaluate whether co-occurrence patterns across protein-coding variants are shared or population-specific across human populations, we apply the statistical framework described above to PHs retained after applying the predefined frequency threshold (≥0.05%) across the five continental super-populations (AFR, AMR, EAS, EUR, and SAS). Analyses are performed both on the aggregated (global analysis) and within each super-population separately (ancestry-stratified analysis). For multiple variants present in each PH, global forest plots summarizing LEC estimates, 95% confidence intervals, and FDR-adjusted significance classifications are generated to quantify enrichment effects, including co-occurrence and mutual exclusivity, alongside Tanghulu-style plots representing variant presence across PHs. Super-population–specific LEC estimates and significance classifications are visualized using ancestry-stratified forest plots displaying global and super-population–specific estimates side by side. Together, these analyses enable direct comparison of multi-variant co-occurrence patterns across populations and reveal both conserved and population-restricted of PH organization.

All analyses are performed in the R programming language (version 4.4.2) using RStudio and on MCW’s Research Computing Cluster.

## Results

### Inter-Individual Genetic Variation in BAF Genes, Individually Missense Intolerant, Are Common

We first catalog the extent and prevalence of genetic variation across the cohort and identify 469 protein-coding variants across 31 BAF-complex genes (**Table.S1**). The majority are moderate impact variants (97.7%, 458/469), whereas only 2.3% (11/469) are of high impact. Missense variants constitute the predominant variant type, accounting for 78.5% (368/469) of all variants, followed by in-frame indels (14.9%, 70/469), comprising 36 insertions and 34 deletions. Normalizing by coding sequence length reveals substantial heterogeneity in variant density across BAF-complex genes. The highest variant densities are observed in *BCL7A* (15.8 variants/kb CDS) and *ARID1B* (11.64), followed by *BICRA* (10.04) and BRD7 (8.69). High variant densities are also observed for *BICRAL* (8.02), *SMARCA2* (7.12), and *ARID1A* (7.00). In contrast, several genes exhibit markedly lower variant burden, including *DPF2* (0.85), *BCL7B* (1.64), *ACTA2* (1.76), and *SMARCB1* (1.73) (**Fig. 1**). Notably, this heterogeneity is not driven by missense intolerance scores as most of the BAF genes studied herein are missense tolerant based on gene-level constraint metrics derived from the Genome Aggregation Database^26^. However, co-occurrence patterns within and between genes may be inflating some missense intolerance statistics. Thus, even among healthy adults, there are common variations within these critical epigenetic regulators, and their patterns are incompletely understood.

**Figure 1.**
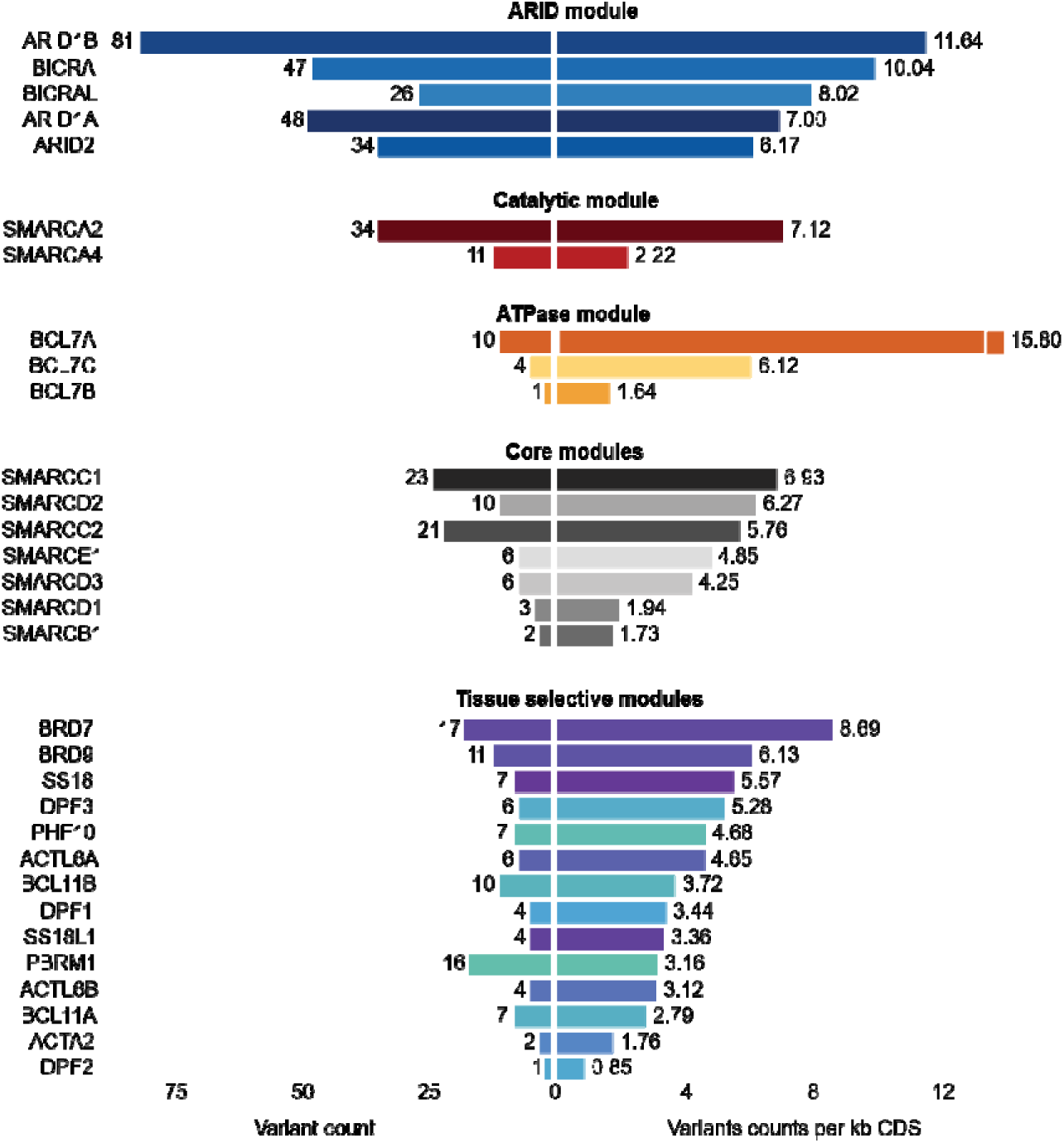
Gene-level distribution of protein-coding variant burden across BAF-complex genes. The left side shows counts of distinct protein-coding variants identified across 31 BAF-complex genes (469 total variants). The right panel shows variant counts normalized by coding sequence length (variants per kb CDS) based on Ensembl MANE Select transcripts. Genes are grouped by functional modules within the BAF complex. Comparison of raw and normalized measures reveals substantial heterogeneity in variant burden that is not fully captured by raw counts alone. Several genes, including *BCL7A*, *BICRA*, *BRD7*, and *BICRAL*, exhibit high normalized variant densities despite relatively modest raw variant counts, indicating disproportionate enrichment after length normalization. In contrast, large genes such as *ARID1A* and *ARID1B* show high raw counts driven in part by gene size but remain enriched after normalization. Genes encoding core structural subunits, including *SMARCB1*, *ACTA2*, *DPF2*, and *BCL7B*, display low normalized variant density, consistent with increased selective constrain. Overall, these patterns indicate that protein-coding variation is unevenly distributed across the BAF complex, preferentially enriched in regulatory and accessory modules of the BAF complex relative to core scaffold components.

### The 1000 Genomes Phase 3 Cohort Exhibits Broad Ancestral Diversity and Balanced Sex Representation

The 1KGP cohort consists of 3,202 individuals distributed across the five continental superpopulations. The largest representation is from the African superpopulation, comprising 27.9% (893/3,202) of individuals, followed by European 19.8% (633/3,202), South Asian, 18.8% (601/3,202), American, 15.3% (490/3,202), and East Asian, 18.3% (585/3,202). Sex distribution is balanced, with 50.1% females (1,603/3,202) and 49.9% males (1,599/3,202), indicating no broad demographic imbalance in the cohort.

### GWAS integration reveals shared SNP–gene–trait architecture

To assess population-level variation in the BAF complex in the context of human disease, we analyze genome-wide association study (GWAS) signals across 31 BAF-complex genes using data from the NHGRI-EBI GWAS Catalog^4^. This analysis identifies 1,343 unique risk alleles associated with 904 traits, which are integrated with 469 curated protein-coding variants from the 1KGP. Applying this proximity-based filtering framework, two genes (*ACTA2* and *ACTL6A*) are excluded due to their roles in numerous actin-dependent enzymes, yielding a refined set of 29 genes with overlapping GWAS and coding variation signals. Within this set, we identify 414 unique alleles across 395 unique protein-coding sites and associated with 416 distinct traits (**Table S2**). Notably, 82 traits are identified independently by SNPs across at least two BAF-complex genes, indicating widespread pleiotropy within the complex (**Table S3**). A representative subset of 16 shared traits highlights gene–trait connectivity mediated by shared SNP–trait associations, based on the most significant SNP (lowest P value) per gene–trait pair with themes enriched for cardiometabolic and hematological phenotypes (**Fig. 2A**). Examples include body mass index, lipids (e.g., total cholesterol measurement, triglyceride measurement, and blood VLDL cholesterol amount), blood pressures (systolic and diastolic blood pressure), and erythrocyte-related measures (e.g., mean corpuscular hemoglobin and erythrocyte volume). Neuropsychiatric traits, including schizophrenia, bipolar disorder, intelligence, and memory performance, are also represented. Recurrent co-occurrence of genes across multiple traits is evident. For example, *ARID1A*, *SMARCA4*, and *SMARCC1* in lipid-related traits, and *BCL7A, PBRM1*, and *SMARCA2* across anthropometric and physiological traits. Certain variants, such as rs114165349 in *ARID1A*, appear across multiple traits and genes, suggesting potential regulatory hotspots underlying pleiotropic effects. Together, these findings define a focused set of BAF-complex loci in which population variation co-localizes with disease-associated signals, pointing to shared biological pathways underlying diverse human conditions.

**Figure 2.**
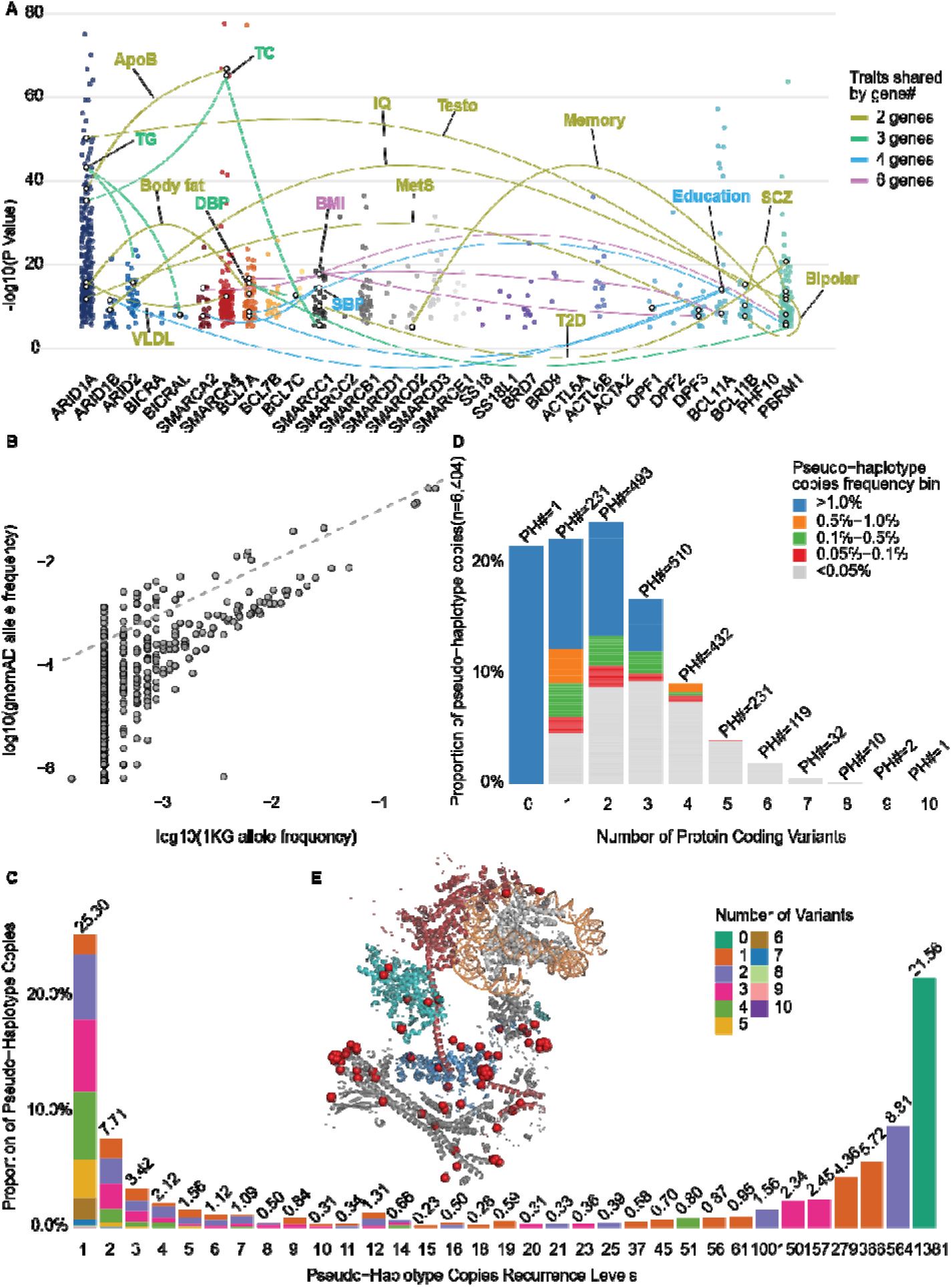
Population variation, pleiotropy, and structural organization of BAF-complex pseudo-haplotypes. **(A)** Visualization of GWAS associations across BAF-complex genes. Each point represents a SNP mapped to a gene–trait pair, with the y-axis showing −log₁₀(P value) and the x-axis indicating gene identity. Highlighted points correspond to the most significant SNP (lowest P value) for each gene–trait pair and are connected by arcs to denote shared trait associations across genes. For a subset of 16 representative traits (from 82 total shared traits, see **Table.S3)** arc and label colors indicate the number of genes sharing each trait. Trait abbreviations: body mass index (BMI), educational attainment (Education), systolic blood pressure (SBP), diastolic blood pressure (DBP), total cholesterol measurement (TC), triglyceride measurement (TG), apolipoprotein B measurement (ApoB), bipolar disorder (Bipolar), blood VLDL cholesterol amount (VLDL), body fat percentage (Body fat), intelligence quotient (IQ), memory performance (Memory), metabolic syndrome (MetS), schizophrenia (SCZ), testosterone measurement (Testo), and type 2 diabetes mellitus (T2D). **(B)** Comparison of variant allele frequencies for the studied variants 1KGP and gnomAD v4.1.0. **(C)** Distribution of PH copies (n = 6,404) across recurrence levels, stratified by protein-coding variant count. The x-axis indicates PH recurrence level (number of identical PH copies), and the y-axis shows the proportion of total PH copies. Stacked bars represent variant count per PH (0–10 variants). The fully reference (0 variants) PH is the most prevalent (1,381 copies; 21.56%). PHs observed at <0.05% frequency (recurrence level <4) are classified as rare and omitted from downstream analyses. **(D)** Distribution of PH copies by variant count, stratified by frequency bins. Bars show the proportion of PH copies grouped by number of protein-coding variants (0–10). Numbers above bars indicate the number of distinct PH configurations (PH#) within each variant-count category. **(E)** Structural mapping of protein-coding variants onto a three-dimensional model of the cBAF complex (based on PDB9A0K^27^). The model includes 10 cBAF subunits (SMARCA4, ARID1A, SMARCC1, SMARCC2, SMARCD1, SMARCE1, ACTB, ACTL6A, SMARCB1, and DPF2), DNA, and histone components. A subset of 118 variants from the total 469 curated protein-coding variants is mapped onto the structure, with the remainder occurring in other configurations of BAF complexes. Of these, 37 variants are localized to resolved residues and shown as red spheres, whereas 81 variants in unresolved regions are represented using pseudo atom placements to approximate their spatial positions. Proteins are displayed as cartoons colored by subunit, with DNA shown in orange and histones in grey. The full distribution of all 469 protein-coding variants across amino acid positions in BAF-complex genes is shown in **Figure. S1C**.

### Allele frequencies are concordant between 1KGP and gnomAD

To validate allele frequency estimates, we compare frequencies derived from 1KGP with those reported in gnomAD v4.1.0 (**Figure.2B**). Of the 469 variants analyzed, 425 are successfully matched across datasets. Allele frequencies show a significant positive correlation (R² = 0.522, p < 0.001), demonstrating strong concordance between 1KGP-derived estimates and large-scale population reference data. This agreement supports the reliability of our current frequency estimates, and indicates the importance of their functional interpretation, while acknowledging that additional diversity may emerge with expanded sampling.

### BAF-Complex Pseudo-Haplotypes Exhibit Extensive Multi-Variant Burden and a Structured Recurrence Landscape

Across 3,202 individuals, 6,404 pseudo-haplotype (PH) copies are constructed from 31 BAF-complex genes, spanning 469 curated protein-altering variant loci and forming 2,062 pseudo-haplotype configurations (Table.S4) (see Methods: Pseudo-Haplotype Construction and Frequency-based Filtering)

At the PH configuration level, we next examine the distribution of protein-coding variant burden across PHs (**Figure.2C–D, Tables. 3–4**). The PH landscape is strongly skewed toward low-frequency configurations (**Figure 2C**, **Table 3**). PHs observed at recurrence level 1 (singletons) account for 25.30% of all 6,404 PH copies and span a wide range of variant counts (1–10 variants), with peak contributions from PHs carrying 3–4 variants. As recurrence level increases, both the diversity and burden of variants decline. PHs at recurrence level 2 predominantly carry 2–3 variants (range: 1–7), while recurrence level 3 PHs are largely restricted to 1–3 variants (range: 1–5). By recurrence level ≥4, PHs are increasingly dominated by low-burden configurations (1–2 variants), and PHs carrying ≥4 variants become progressively rare. Overall, higher variant burdens are largely confined to low-recurrence PHs, suggesting selective or structural constraint against accumulation of multiple coding variants within recurrent configurations. Consistent with this pattern, a small number of highly recurrent PHs account for a substantial fraction of all PH copies. The full reference (0-variant) configuration is the most abundant, observed 1,381 times (21.56%). Additional recurrent PHs occur at frequencies of 564 (8.81%), 366 (5.72%), and 279 (4.36%), demonstrating a strongly right-skewed recurrence distribution dominated by a few common configurations.

**Table 3.**
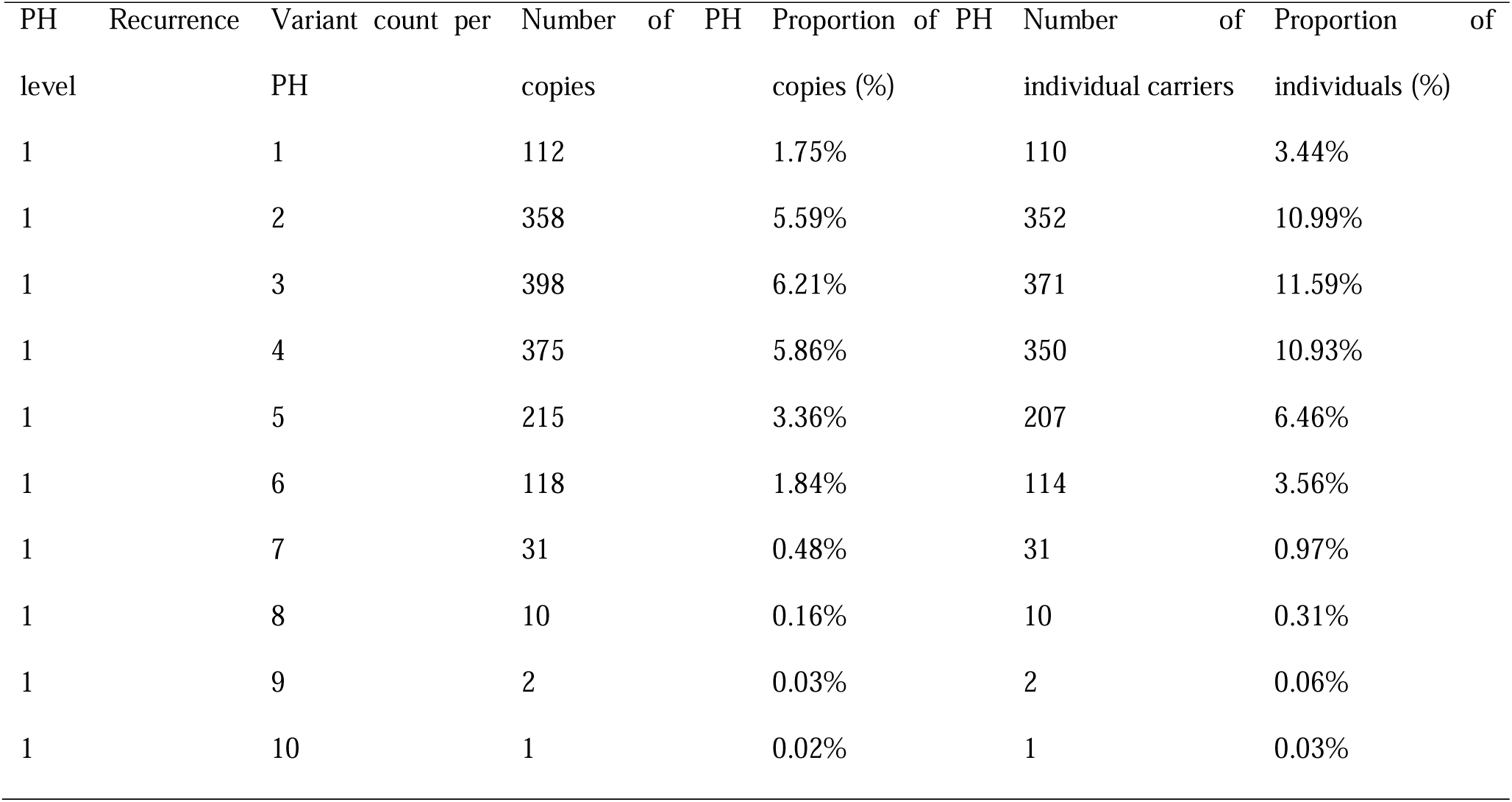
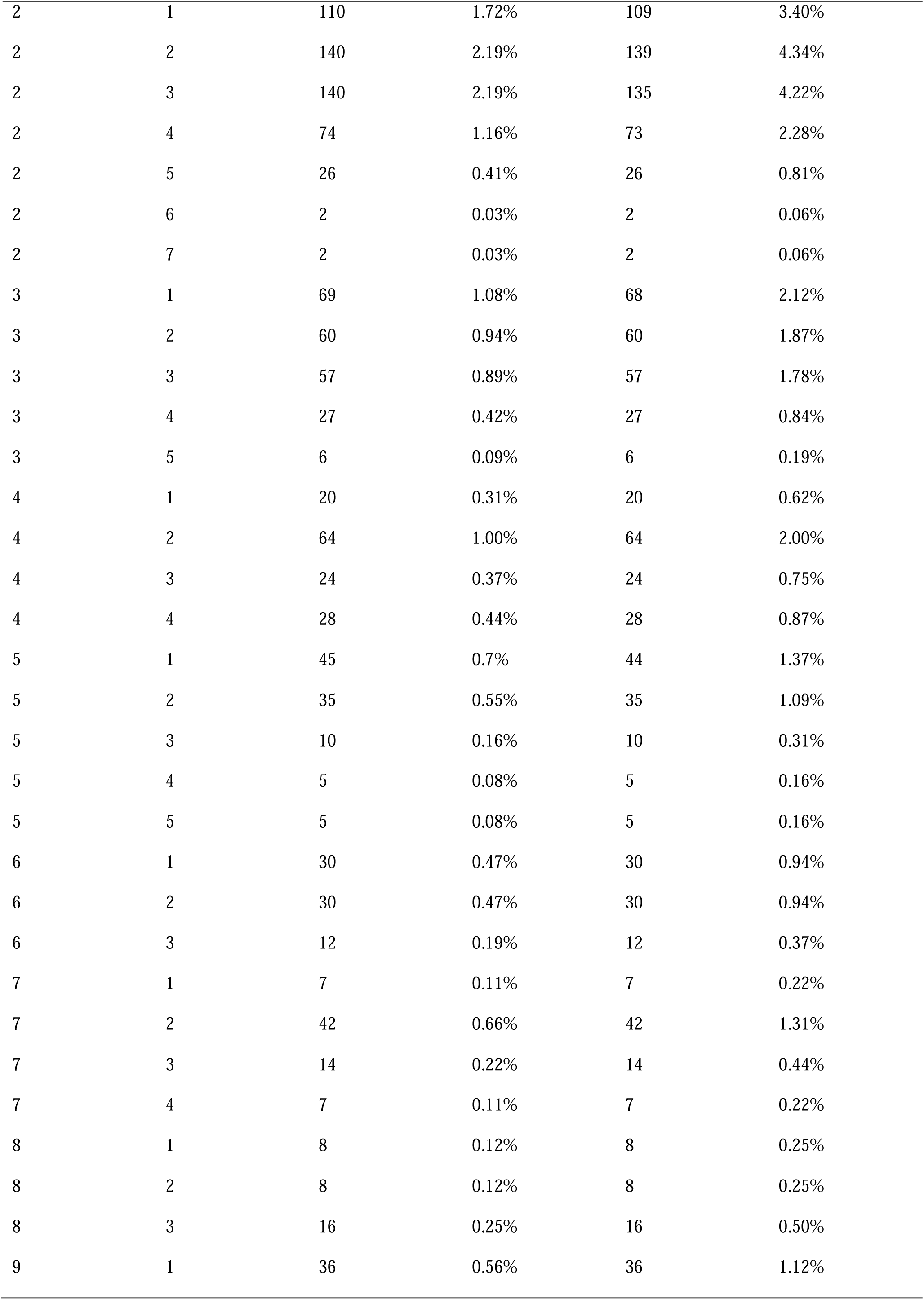
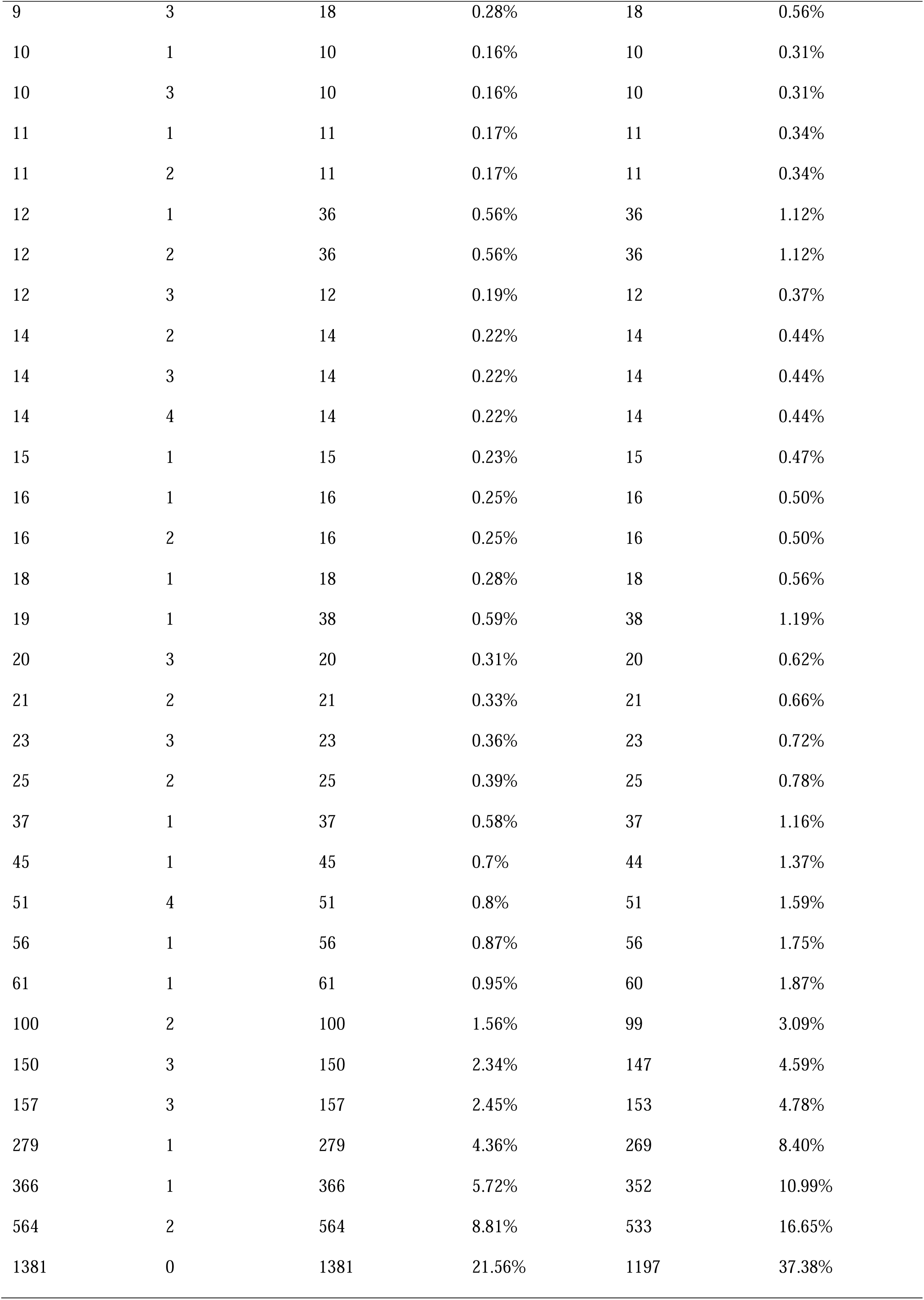
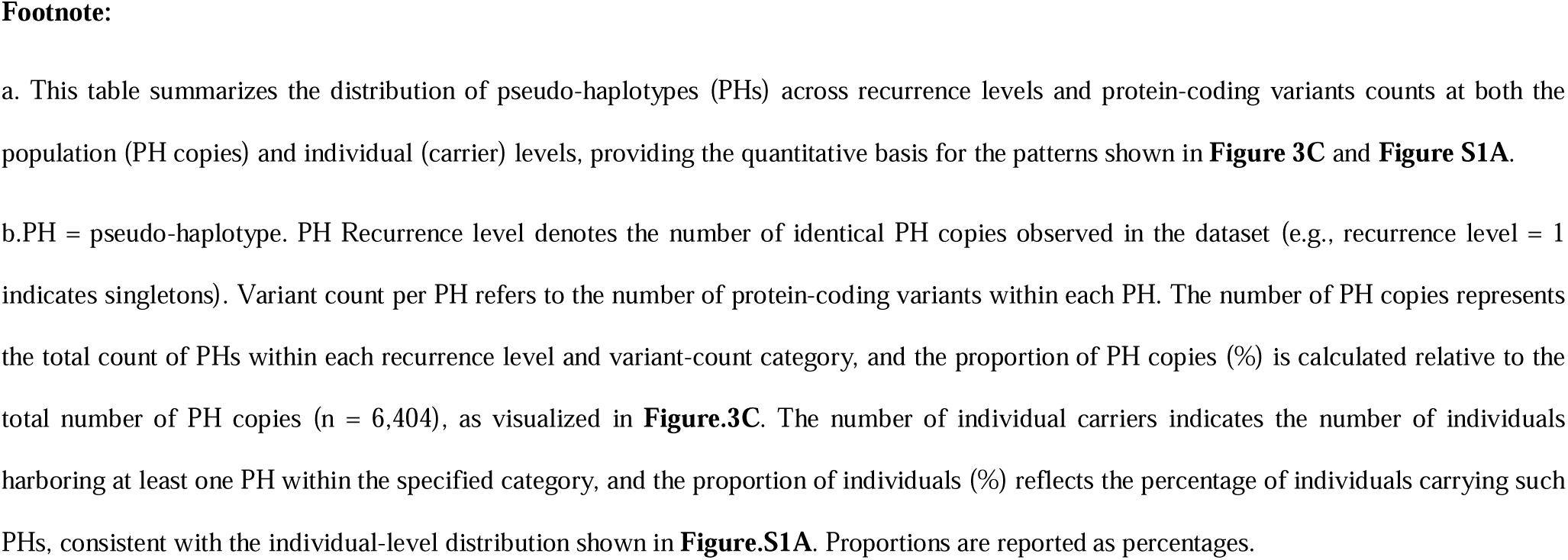
Distribution of PH copies and individual carriers by recurrence level and variant count.

Analysis by variant number further confirms that low-burden PHs dominate the population (**Fig. 2D**, **Table 4**). PHs carrying 0–3 variants comprise the majority of all PH copies (0 variants: 21.56%; 1 variant: 22.25%; 2 variants: 23.80%; 3 variants: 16.79%) and are enriched in higher-frequency bins. In contrast, PHs with ≥4 variants decline sharply in prevalence (4 variants: 9.07%; 5 variants: 3.94%; ≥6 variants: ≤1.87% each) and are primarily restricted to low-frequency or rare categories.

**Table 4.**
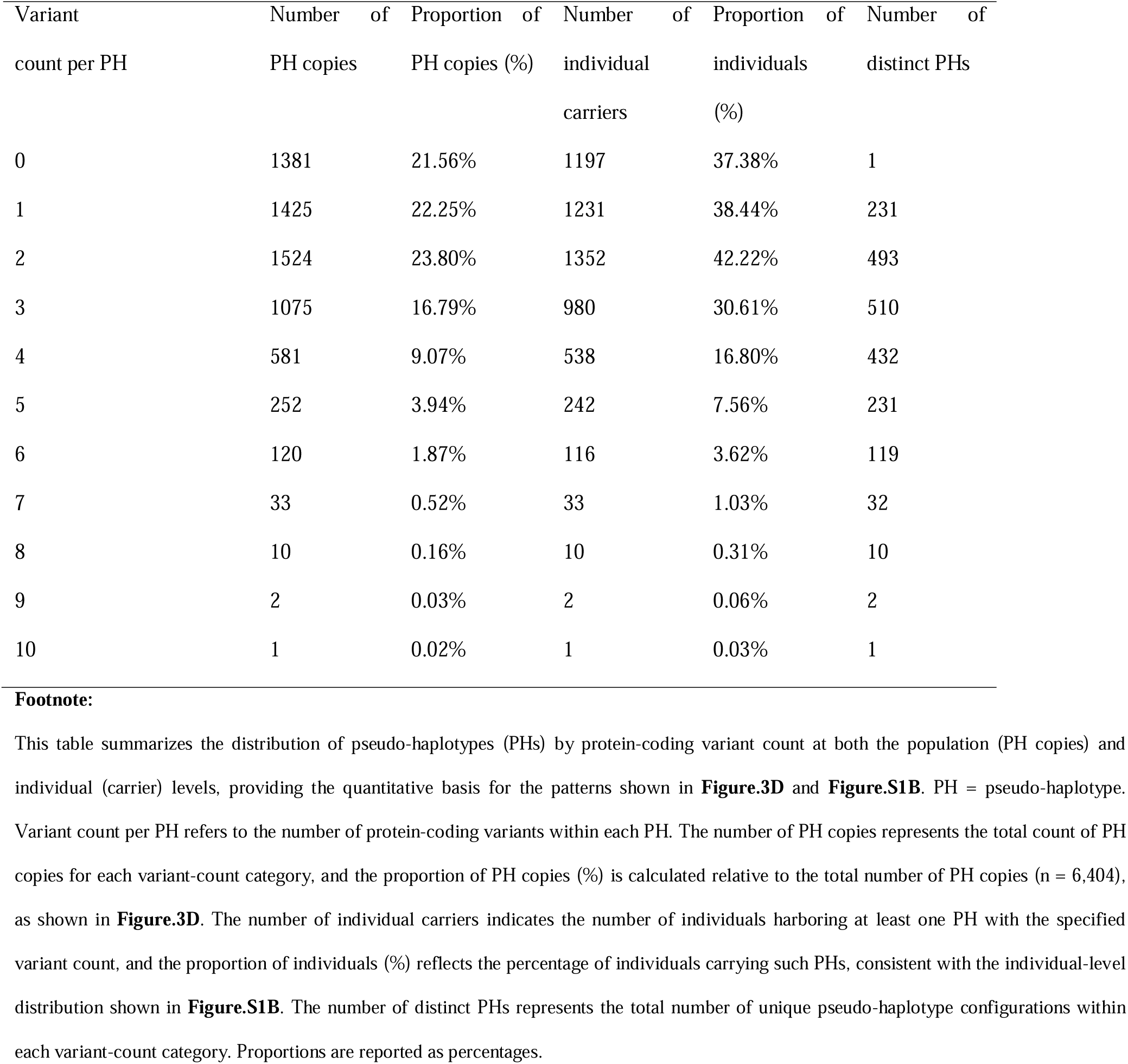
Distribution of PH copies and individual carriers by variant count.

Overall, 78.44% of PH copies carry at least one coding variant, 56.18% carry ≥2 variants, and 32.39% carry ≥3 variants, spanning a range of 1–10 variants per PH. To reduce instability from low-recurrence configurations, PHs below the predefined PH frequency threshold (≥0.05%) are excluded from downstream analyses. The retained PHs are subsequently used to identify recurrent multi-variant configurations, including combinations involving *BICRA* and *SMARCA2*.

At the individual level, a concordant distribution of PH recurrence and variant burden was observed (**Fig. S1A–B, Tables 3–4**). **Figure S1A** summarizes recurrence-level distributions stratified by variant burden, whereas **Figure S1B** shows the distribution of protein-coding variant burden per PH at the individual level. Individuals carrying low-burden PHs predominated within the population. The reference (0-variant) configuration is present in 37.38% of individuals, while PHs with 1 and 2 variants are observed in 38.44% and 42.22% of individuals, respectively (**Fig. S1B, Table 4)**. Carrier frequency declines with increasing variant burden, with PHs carrying 3 variants present in 30.61% of individuals and higher-burden configurations occurring less frequently (4 variants: 16.80%; 5 variants: 7.56%; ≥6 variants: ≤3.62%). Despite this decline, multi-variant configurations remain common in the population.

Collectively, these results reveal a structured recurrence landscape characterized by (i) dominance of low-burden PHs, (ii) strong constraint against high variant burden, and (iii) a small number of highly recurrent configurations that capture a large proportion of population variation. This structure supports robust downstream co-occurrence analyses and highlights strong functional constraint across the BAF complex in human populations.

### Structural mapping reveals spatial clustering of coding variation

To investigate the structural distribution of coding variation, we map a representative subset of protein-coding variants of 469 variants onto a three-dimensional model of the canonical BAF (cBAF) complex (**Fig. 2E**). This model incorporates 10 core cBAF subunits with resolved structural information. From the total set of 469 curated protein-coding variants, 118 variants are selected for structural visualization based on their map ability to resolve or approximated regions of the complex. Mapped variants display a non-random spatial distribution across the complex. While many variants are dispersed throughout peripheral or structurally flexible regions, a subset forms localized clusters within specific subunits and structural interfaces, particularly within ARID1A, SMARCA4, and SMARCC1. These clusters are frequently observed near regions involved in protein–protein interactions or DNA engagement, suggesting potential hotspots of functional sensitivity. In summary, these observations indicate that protein-coding variation within the BAF complex is both structurally constrained and spatially organized, with clustering patterns that highlight potentially important functional regions of the cBAF architecture.

### BAF-Complex Pseudo-Haplotypes Exhibit Structured Multi-Variant Diversity and Recurrent Variant Backbones

At the predefined 0.05% PH frequency threshold, we identified 122 distinct PHs representing recurrent protein-coding variant configurations across the cohort (**Table S4**). Consistent with the global distribution described above (**Figure 2C–D**; **Tables 3–4**), these recurrent PHs predominantly comprise low- to moderate-burden configurations (0–5 variants per PH) and span a broad frequency range. Most PHs are defined by multi-gene variant combinations across BAF-complex genes, including BICRA, SMARCA2, BCL7A, ARID1A, ARID1B, ARID2, SMARCA4, BRD7, BRD9, ACTL6A, SS18, SS18L1, PBRM1, DPF3, BICRAL, SMARCC2, and SMARCD3. Protein-coding variants exhibit strong, non-random clustering across PHs, with recurrent co-occurrence of BICRA (e.g., p.Pro683Ser and p.Thr1044Ala) and SMARCA2 variants (e.g., p.Gln238del and p.Asp1546Glu) forming a dominant backbone of multi-variant configurations (**Fig. 3A**). In contrast, lower-frequency PHs incorporate additional variants—particularly in-frame indels in ARID1A, ARID1B, and ARID2—resulting in increasingly complex configurations that segregate into distinct branches within the PH similarity tree according to shared variant architectures. These observations indicate a hierarchical organization of pseudo-haplotype diversity, in which a limited number of recurrent or founder variants define the basal pattern while additional variants accumulate thereupon to generate less frequent configurations (**Fig. 3B**). Overall, these patterns indicate that BAF PH diversity is dominated by a modest number of widely shared variant combinations, whereas a long tail of lower-frequency PHs contributes substantial combinatorial complexity and inter-individual differences.

**Figure 3.**
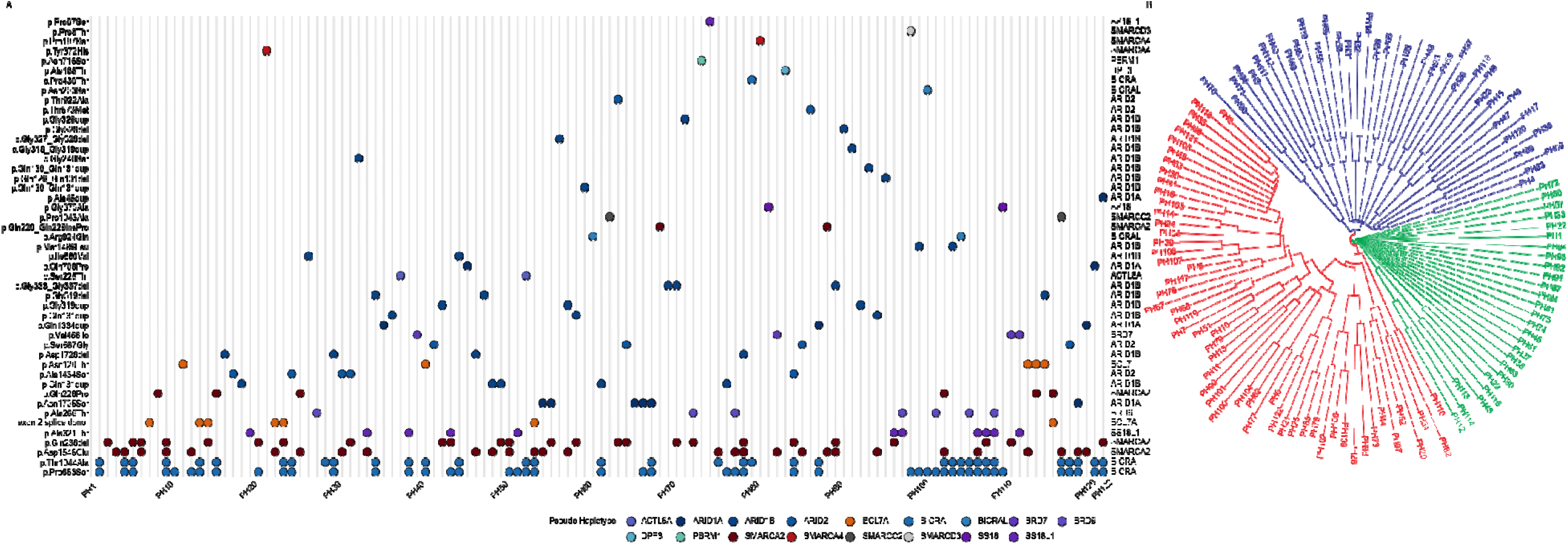
Co-occurrence patterns and similarity structure of pseudo-haplotypes at the 0.05% frequency threshold. **(A)** Tanghulu-style co-occurrence plot of protein-coding variants across the 122 PHs retained at the 0.05% PH frequency threshold (**Table S4**). Columns represent PHs, and dots indicate the presence of protein-coding variants, colored by gene. A prominent co-occurrence module centered on BICRA p. Pro683Ser, frequently accompanied by BICRA p. Thr1044Ala and recurrent SMARCA2 variants (e.g., p. Gln238del, p. Gln228Pro, p. Asp1546Glu), forms a dense vertical band consistent with recurrent co-inheritance of a core variant set. In contrast, ARID1A/ARID1B/ARID2 indels, particularly in the ARID1B glycine-rich region alterations (e.g., p. Gly319dup/del, p. Gly333_Gly337del, and nearly glycine-repeat expansions), occur more sporadically and are enriched among lower-frequency pseudo-haplotypes. Variants in SS18L1, BRD9, and BCL7A appear as secondary modifiers of the predominant combinations. PH identifiers (PH#) are ordered from lowest to highest PH copy frequency, consistent with **Table S4**; x-axis labels are displayed at intervals of 10 PHs for visualization clarity. **(B)** Circular PH similarity tree constructed using the Jaccard index to quantify pairwise similarity in variant composition. Branches segregate into three major color-coded clusters based on shared variant architecture. Cluster I (green) comprise heterogeneous pseudo-haplotypes enriched for ARID1B, ARID1A, and other SWI/SNF structural subunit variants, often defined by unique or low-frequency combinations and lacking a recurrent core variant set. Cluster II (red) forms the largest clade and is characterized by a recurrent BICRA–SMARCA2 variant backbone, most commonly involving BICRA p.Pro683Ser) with *BICRA* p. Thr1044Ala and SMARCA2 alterations, from which numerous pseudo-haplotypes radiate through stepwise acquisition of additional variants (e.g., SS18L1, BRD9, ARID1B). Cluster III (blue) lacks the recurrent BICRA p.Pro683Ser backbone and is instead driven primarily by SMARCA2 (e.g., p. Gln238del, p. Gln228Pro, p. Asp1546Glu) affecting ATPase/helicase -associated regions, with secondary contributions from *ARID* family genes and *BRD7*. Major internal branches reflect shared backbone architectures, whereas long terminal branches represent PHs defined by rare or low-frequency coding events.

### Global Co-Occurrence Analysis Reveals Strong Multi-Variant Enrichment and a Dominant BICRA–SMARCA2 Backbone

Using the LEC-based multi-variant enrichment framework described (see **Methods***: Statistical Analysis of Multi-variant Enrichment*), we evaluated combinatorial interactions among protein-coding variants across PHs retained at the predefined 0.05% PH frequency threshold and containing at least two alternative alleles. Analyses were performed in both the aggregated cohort and ancestry-stratified super-populations to assess the consistency of co-occurrence patterns across populations.

Across the 122 retained PHs, 80 contain at least two protein-coding variants and are evaluated for multi-variant enrichment (**Table S5**). Among these, 57 PHs show significant deviation from independence (FDR < 0.05), with the vast majority exhibiting positive enrichment, indicating widespread co-occurrence of protein-coding variants (**Fig. 4A-B**). The strongest and most consistent enrichment signals are observed among PHs containing BICRA and SMARCA2 variants. Pairwise combinations such as PH2 (BICRA p.Pro683Ser and p.Thr1044Ala) show robust enrichment (LEC= 1.72, 95% CI 1.65–1.80, FDR < 0.001). Higher-order configurations, such as PH5, PH6, PH10 incorporating SMARCA2 variants (e.g., p.Gln238del, p.Asp1546Glu) further increase enrichment magnitude, with LEC values frequently exceeding 1.4 and up to ∼3.2 (FDR < 0.001), consistent with strong multi-variant aggregation. While most PHs exhibit modestly positive enrichment values, these BICRA–SMARCA2 combinations consistently display the largest effect sizes, indicating substantial and reproducible co-occurrence relative to expectations under independence. In contrast, evidence of mutual exclusivity is limited to a small number of PHs like PH26 (SMARCA2 p.Gln228Pro with p.Asp1546Glu) which show modest depletion (LEC= −0.72, 95% CI −1.40 to −0.08, FDR = 0.025). Overall, positive co-occurrences signals dominate, indicating that protein-coding variants tend to aggregate non-randomly within PHs rather than segregate independently, supporting a structured and biologically meaningful organization of variant co-occurrence at the population level.

**Figure 4.**
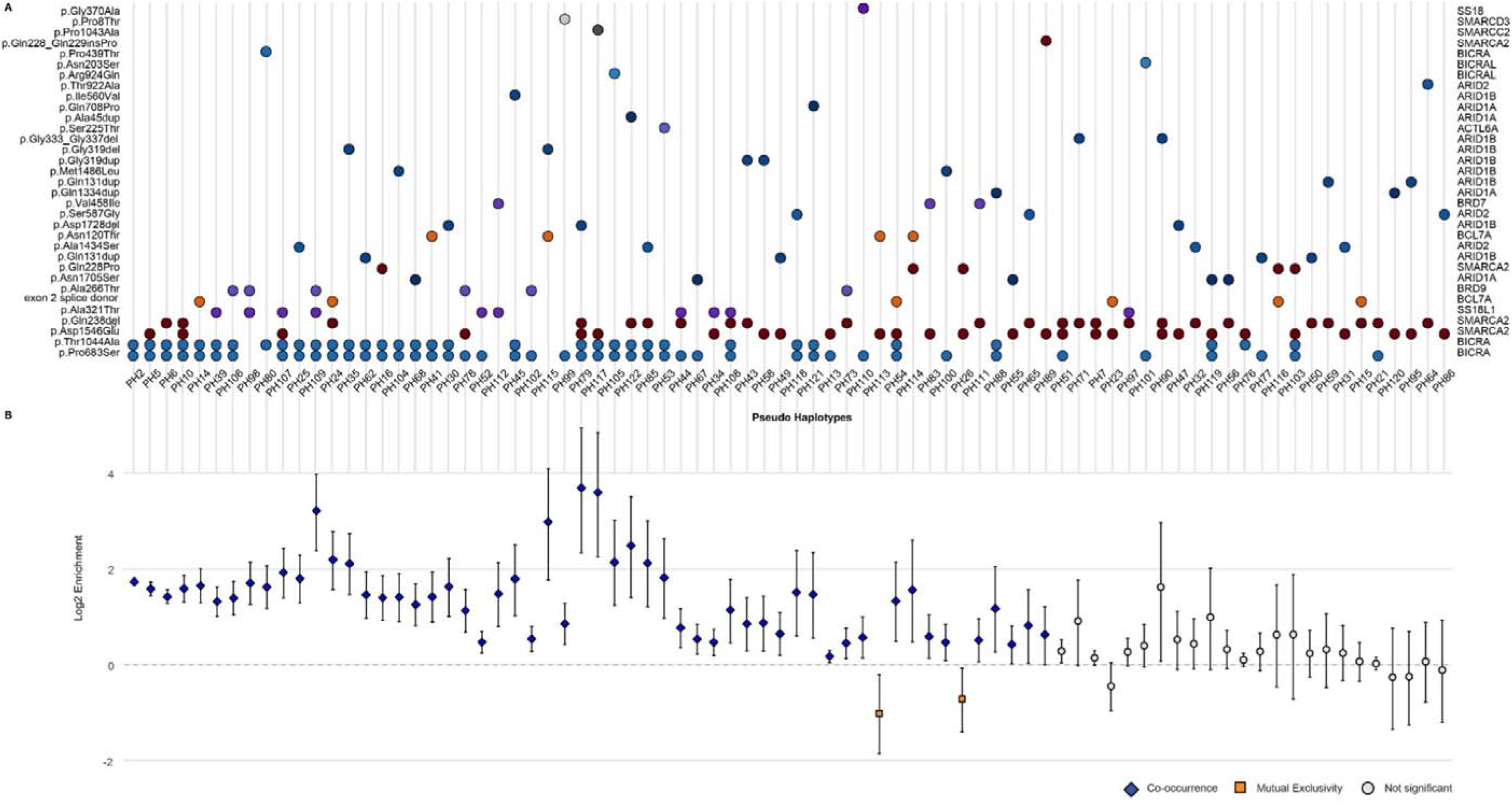
Co-occurrence associations and variant composition of pseudo-haplotypes at the 0.05% frequency threshold. **(A)** Variant composition matrix displaying the presence (colored dots) or absence of each protein coding variant along the vertical axis, with columns representing the 80 PHs that contain ≥2 variants and pass the 0.05% **frequency** threshold (**Table S5**). Dots are colored by gene. Dense clusters highlight frequent PHs dominated by BICRA and SMARCA2 variants, whereas ARID1A, ARID1B, ARID2, and BCL7A variants appear more sparsely and characterize lower-frequency, often lineage-specific haplotypes. Pseudo-haplotype identifiers (PH#) are ordered from highest to lowest copy frequency, consistent with **Table S4** and Figure 3A. **(B)** Forest plot summarizing the enrichment strength (log_2_ Enrichment) of protein coding variant combinations for these PHs (**Table S5**). Each point represents the highest-order variant combination within a PH (i.e., the full multi-variant set observed in that PH), with error bars indicating the 95% confidence interval of the log_2_ Enrichment. Blue diamonds denote statistically significant co-occurrence (positive log_2_ enrichment values with FDR < 0.05), orange squares indicate significant mutual exclusivity (negative log_2_ enrichment values with FDR < 0.05), and grey circles represent non-significant associations (FDR ≥ 0.05). PHs are ordered from lowest to highest FDR (left to right), corresponding to decreasing statistical significance.

### Super-Population Co-Occurrence Patterns Reveal Conserved and Ancestry-Specific Variant Interactions

To assess whether multi-variant co-occurrence patterns are conserved across genetic ancestries, we evaluated LEC within each of the five super-populations (AFR, AMR, EAS, EUR, and SAS) and the aggregated cohort (**Fig. 5, Table S5**). While many globally significant PHs remain enriched across multiple populations, both the magnitude and statistical significance of enrichment vary across ancestries. The most consistent signal is observed for the core BICRA haplotype, PH2 (BICRA p.Pro683Ser and p.Thr1044Ala), which exhibits significant positive enrichment across all five super-populations and in the aggregated cohort (LEC = 1.72, 95% CI 1.65–1.80, FDR < 0.001), indicating a broadly conserved co-occurrence pattern across human populations, including European (2.30, 95% CI 2.12–2.49, FDR < 0.001), East Asian (2.01, 1.83–2.19, FDR < 0.001), American (1.97, 1.77–2.18, FDR < 0.001), South Asian (2.04, 1.86–2.21, FDR < 0.001), and African populations (0.90, 0.75–1.06, FDR < 0.001). Next, PHs defined by BICRA p.Pro683Ser and p.Thr1044Ala and incorporating SMARCA2 variants also show widespread enrichment, but with variable ancestry-specific strength. For example, PH5 (including SMARCA2:p.Asp1546Glu) and PH6 (including SMARCA2:p.Gln238del) are significantly enriched in the aggregated cohort (PH5: 1.58, 1.44–1.72, FDR < 0.001; PH6: 1.41, 1.27–1.56, FDR < 0.001) and across all populations, although the magnitude of enrichment is reduced in African ancestry. In contrast, PH10 (incorporating both SMARCA2 variants) shows strong enrichment in several populations, including European (2.16, 0.61–3.53, FDR = 0.036), East Asian (1.58, 1.03–2.11, FDR < 0.001), American (2.29, 1.50–3.05, FDR < 0.001), and South Asian populations (1.99, 1.32–2.60, FDR < 0.001), but is not significant in African ancestry (0.50, 0.00–0.98, FDR = 0.147), indicating population-dependent differences in co-occurrence strength. Some PHs exhibit population-specific enrichment despite weak or non-significant signals in the aggregated cohort. For instance, PH7 (SMARCA2:p.Gln238del and p.Asp1546Glu) shows only a weak global association (LEC = 0.14, 95% CI −0.01–0.29, FDR = 0.072), but modest enrichment in the American population (0.45, 0.01–0.89, FDR = 0.049). Conversely, certain PHs are observed only in a subset of populations, limiting statistical power due to low counts and resulting in wider confidence intervals. For example, PH24 shows significant enrichment in East Asian (2.75, 1.97–3.51, FDR < 0.001) and South Asian populations (2.65, 1.46–3.81, FDR = 0.001), and global (2.19, 1.57–2.77, FDR < 0.001), but is not observed or not significant in other populations.

**Figure 5.**
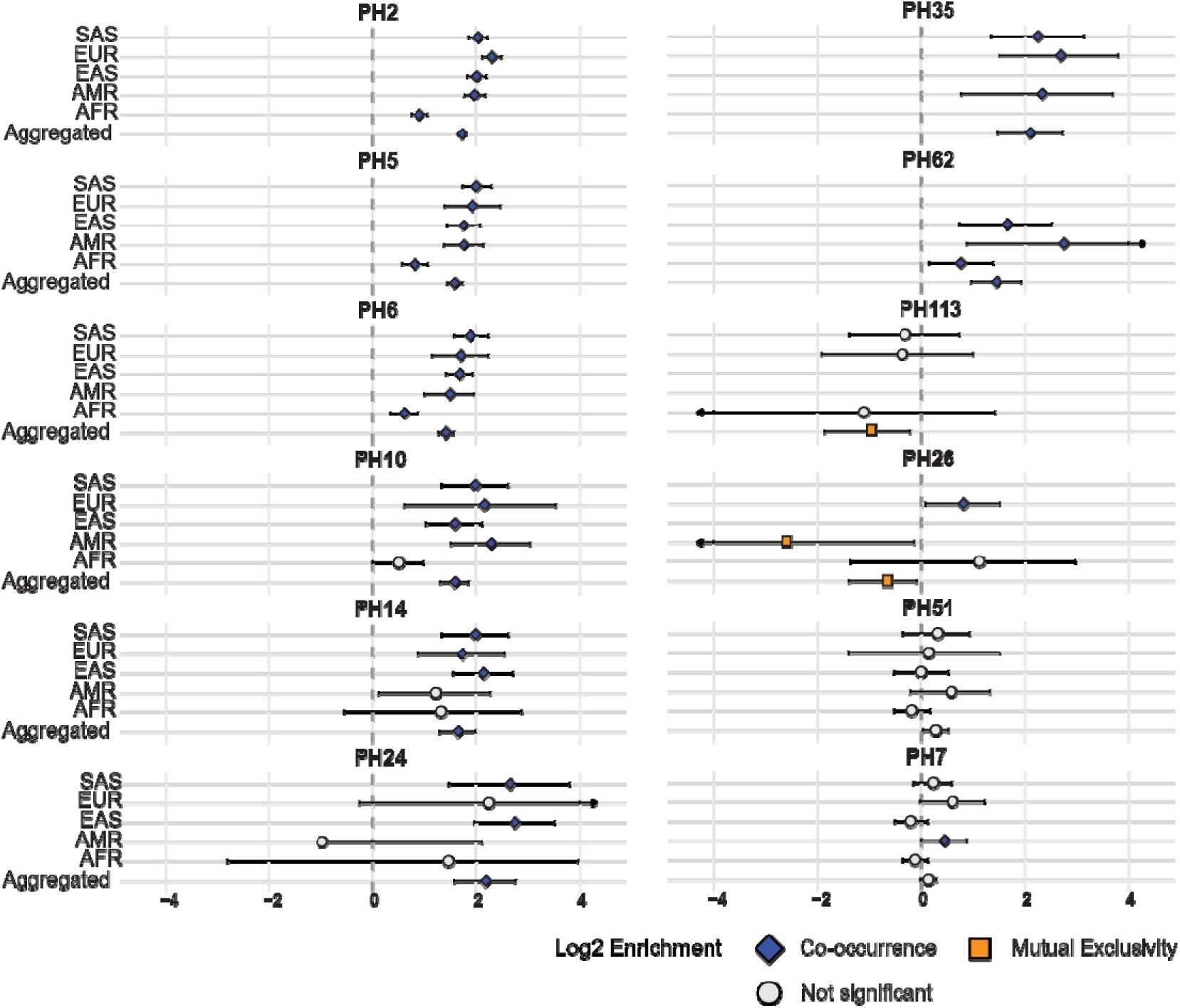
Super-population–stratified co-occurrence patterns of selected PHs at the 0.05% HP frequency threshold. Forest plots showing log_2_ Enrichment (LEC) estimates for representative PHs across the aggregated cohort and five 1KGP super-populations. Each panel corresponds to a single PH, and points represent LEC values with 95% confidence intervals for the full multi-variant configuration defining that PH. The x-axis represents LEC with horizontal error bars indicating 95% confidence intervals. Super-populations are displayed along the y-axis in a consistent order: South Asian (SAS), European (EUR), East Asian (EAS), American (AMR), African (AFR), and the aggregated. Corresponding statistical results, including LEC estimates, confidence intervals, and FDR-adjusted P-values for all PH–population combinations, are provided in **Table.S5**. Aggregated cohort enriched PHs—particularly those defined by recurrent BICRA–SMARCA2 variant combinations (e.g., PH2, PH5, PH6, PH10)—show consistent positive enrichment across multiple super-populations, although the magnitude and significance vary by ancestry. In contrast, some PHs display population-specific signals (e.g., PH24, PH7), while others exhibit depletion or mutual exclusivity in selected populations (e.g., PH26, PH113).

A small number of PHs exhibit negative enrichment consistent with mutual exclusivity. Notably, PH26 (SMARCA2 p.Gln228Pro and p.Asp1546Glu) shows significant negative enrichment in the aggregated cohort (LEC = −0.72, 95% CI −1.40 to −0.08, FDR = 0.025) and in the American population (−2.66, CI −6.92 to −0.13, FDR = 0.026), but not in other populations. Similarly, PH113 shows significant depletion only in the aggregated cohort (−1.02, CI −1.86 to −0.21, FDR = 0.010), indicating context-specific mutual exclusivity. Taken together, these results indicate that pseudo-haplotype structure within the BICRA–SMARCA2 chromatin-remodeling gene network reflects a combination of globally conserved and ancestry-specific co-occurrence patterns. A core set of BICRA-centered haplotypes shows consistent enrichment across ancestries, whereas additional configurations display variability in strength, frequency, and detectability across populations, highlighting the structured and population-dependent nature of multi-variant interactions.

## Discussion

As genomics moves beyond single-gene paradigms, it’s becoming clear that individualized genetics isn’t only about rare variants. Rather, ultra-rare and private variants superimpose on rare combinations of common variants to produce a wide range of inter-individual differences, a phenomenon increasingly recognized in studies of complex trait architecture and polygenic inheritance^28,29^. Accounting for these private and rare constellations can reveal hidden mechanisms, refine risk prediction, and bridge the gap between genome-scale association and patient-specific biology.

Our analysis reveals that 94.3% of 1KGP individuals carry at least one coding sequence variant across the 31 BAF complex genes, while 77.5% carry ≥2 variants and 50.9% carry ≥3 variants across their combined pseudo-haplotype genotypes. This striking demonstration of extensive genetic diversity within this essential chromatin remodeling complex is further supported by the observation that 25.30% of PHs are private (observed in a single individual) within the cohort. This finding is compelling given the BAF complex’s central role in transcriptional regulation and its frequent implication in neurodevelopmental disorders and cancers^30^. The high prevalence of sequence divergence underscores a limitation of the monogenic paradigm for variant interpretation and emphasizes the need for individualized genomic interpretation frameworks^31^. The non-random co-occurrence of variants in, for example, BICRA and SMARCA2 across world populations suggest these genes may act as context-specific modifiers, with coordinated inheritance patterns implying functional epistasis - a phenomenon where combinatorial variant effects diverge from individual allele impacts^32^. Such interactions align with evidence that paralogous BAF subunits exhibit compensatory relationships^7^, where specific variant combinations could buffer or amplify certain phenotypic consequences, that may depend on regional environmental or social pressures on chromatin remodeling pathways, or founder effects shaping PH structures.

The near ubiquity of multi-variant BAF haplotypes provides a mechanistic lens through which to view variable expressivity in associated genetic disorders. For instance, SMARCB1 variants drive phenotypes ranging from benign tumors to aggressive cancers^33^, while ACTL6A mutations exhibit spectrum-specific impacts on neuronal development^34^. Our observations suggest that background genetic architecture could in fact modulate pathogenic variant effects. PH-specific epistasis could explain why identical rare pathogenic variants yield different clinical severities for different individual patients, while population-specific modifier variants may buffer or exacerbate phenotypes expressivity, complicating cross-ancestry clinical predictions^32^. Challenging as they may be to accomplish, these findings advocate for an integrated variant assessment model that moves beyond single-gene and site-specific pathogenicity evaluations to consider multi-locus haplotype effects, local haplotype architectures rather than continental ancestry proxies, topological features of the encoded gene products *in situ*, and three-dimensional chromatin interaction maps, rather than isolated studies^29^.

Further, the field continues to grapple with the complex relationships among incomplete penetrance, variable expressivity, and inter-individual variation in traits^35^ and drug responses, to name a few – a subset of which may be explained by epigenetic PHs that collectively shape individual person’s responses to the same environmental exposures. Because enzymes like BAF regulate the whole genome, epigenetic PHs, which could be termed epi-types, have the potential to modulate any human trait. Other factors intersect with PHs including modifier genes, epigenetic modifications, chromatin variants, environmental influences, genetic predispositions, and the specific location and consequence of genetic variants within gene products^30^. These factors collectively contribute to the mechanistic complexity of genotype-phenotype relationships, presenting significant challenges for improving genetic interpretation and clinical prediction, which we are confident will be addressed through our future studies.

The current study demonstrates that common BAF variants exist on a continuum with rare pathogenic mutations, challenging traditional dichotomous distinctions between benign polymorphisms and pathogenic alleles for variant annotation and interpretation. Both are heterogeneous, superimposed uniquely for many individuals. As global sequencing efforts expand, the observed PH landscape will likely increase, revealing deeper layers of inter-individual BAF complexity. Clinically, this evolution demands haplotype-aware diagnostics to address the limitations of current guidelines that assess variants in isolation, ancestry-calibrated risk scores informed by population-specific haplotype frequencies, and functional atlases mapping pseudo-haplotype effects on chromatin accessibility and transcriptional dynamics. In fact, we anticipate that findings such as these could underly components of existing polygenic risk scores. The BAF complex serves as a paradigm for understanding how layered genetic variation-from common SNPs to rare CNVs-converges on phenotypic outcomes.

Despite these advances, there are two major limitations that must be considered. First, co-occurrence inference was conducted independently within each 1KGP super-population (AFR, AMR, EAS, EUR, SAS), thereby minimizing the risk of spurious signals from population stratification. In contrast, descriptive summaries of pooled (global) prevalence and the overall PH landscape are derived from the aggregated cohort. Pooled analyses remain susceptible to residual confounding from unmodeled admixture or ancestry gradients (e.g., Simpson’s paradox-like effects), which may subtly influence descriptive global patterns even when stratified testing protects inferential claims. Second, no linkage disequilibrium (LD) pruning is applied, and all variants are treated as independent under the null hypothesis, regardless of their physical proximity on the same chromosome or within the same gene. While this approach is appropriate for an exploratory analysis focused on gene-set combinatorial patterns, it increases the risk that observed co-occurrence signals are driven (at least in part) by local LD rather than true biological epistasis or functional selection. Consequently, significant deviations reflect statistical non-independence in multi-locus allele configurations within the BAF complex rather than direct evidence of causal mechanisms, physical haplotype structure, or long-range interactions beyond simple proximity. These limitations are inherent to the study’s pioneering and descriptive focus on variant burden, recurrence, and pooled PH frequency. They do not undermine the core observation of non-random multi-locus configurations but highlight clear directions for methodological refinement as we look towards application of the PH framework to larger population genomics studies, such as All of Us Research Program^34^. Future work can address them through local ancestry deconvolution, LD pruning (e.g., r²-based block formation), whole-chromosome phasing, expanded LD thresholds, inverse-variance weighted meta-analysis of super-population effect sizes to derive robust global estimates, and functional validation of prioritized configurations.

## Conclusion

This study introduces a comprehensive methodology for PH analysis of BAF complex genes using 1KGP data, enabling exploration of genetic variation and co-occurrence patterns across multiple chromatin remodeling genes that together accomplish this essential cellular function. Our analysis reveals that common PHs exhibit substantial genetic diversity and global hierarchical relationships. Given the central role of the BAF complex in transcriptional regulation, chromatin remodeling, and cellular differentiation, and the implication of its component genes in a wide range of human diseases and individual-level risk prediction and interpretation in sequencing studies. This study provides a valuable framework for investigating how genetic diversity and haplotype structure within the BAF complex may contribute to phenotypic diversity and disease susceptibility, and of PHs in general. Overall, our findings show that PH-based analysis is a valuable tool for uncovering genetic organization and diversity of key chromatin remodeling genes like those in the BAF complex. Future research can extend this approach to other complexes and cellular processes, incorporate finer-scale ancestry and LD controls, inverse-variance weighted meta-analysis for cross-population synthesis, and evaluating the clinical and trait-based consequences of multi-gene PHs. We envision PH analysis as a component of defining mechanistic links among human diseases according to their underlying molecular effects on protein complexes and cellular architectures.

## Acknowledgments

Research reported in this publication was supported by the National Institute of General Medical Sciences of the National Institutes of Health under Award Number R35GM153740. The content is solely the responsibility of the authors and does not necessarily represent the official views of the National Institutes of Health. This project was also funded in part by the Advancing a Healthier Wisconsin Endowment at the Medical College of Wisconsin and with computational resources and technical support provided by the Research Computing Center at the Medical College of Wisconsin.

## Author contribution statement

Conception and design: MZ, XD. Development of methodology: XD, MZ. Acquisition of data: XD, MZ. Analysis of data: XD. Interpretation of data: XD, NH, JW, MZ. Writing and revising of the manuscript: XD, MZ. Review of the manuscript: XD, NH, JW, MZ.

## Data and code availability

Source data are provided in **Table 1**. The code supporting this study is available from the corresponding author upon reasonable request.

## Declaration of generative AI and AI-assisted technologies in the writing process

During the preparation of this work, the authors used ChatGPT to assist with language refinement, and formatting of the manuscript. After using this tool, the authors critically reviewed and edited the content and take full responsibility for the final version.

