## Supplementary figures and images for "Defining Pseudo-Haplotype Analysis Reveals Multi-Gene Genetic Pattern Across BAF Chromatin Remodeling Complexes"

### Supplementary Figure 1A-C

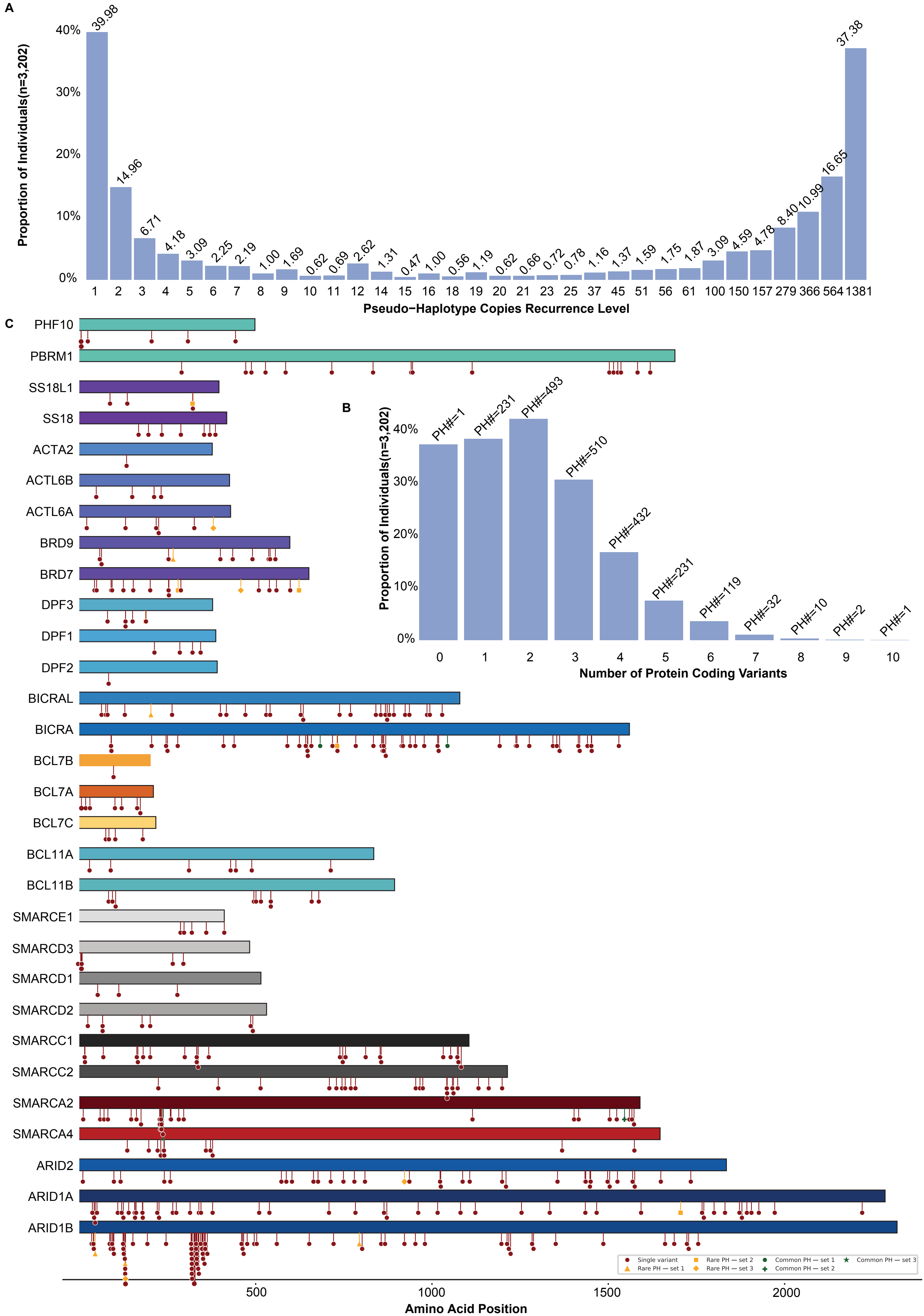
