## Supplementary material for "Defining Pseudo-Haplotype Analysis Reveals Multi-Gene Genetic Pattern Across BAF Chromatin Remodeling Complexes"

### Supplemental Figures

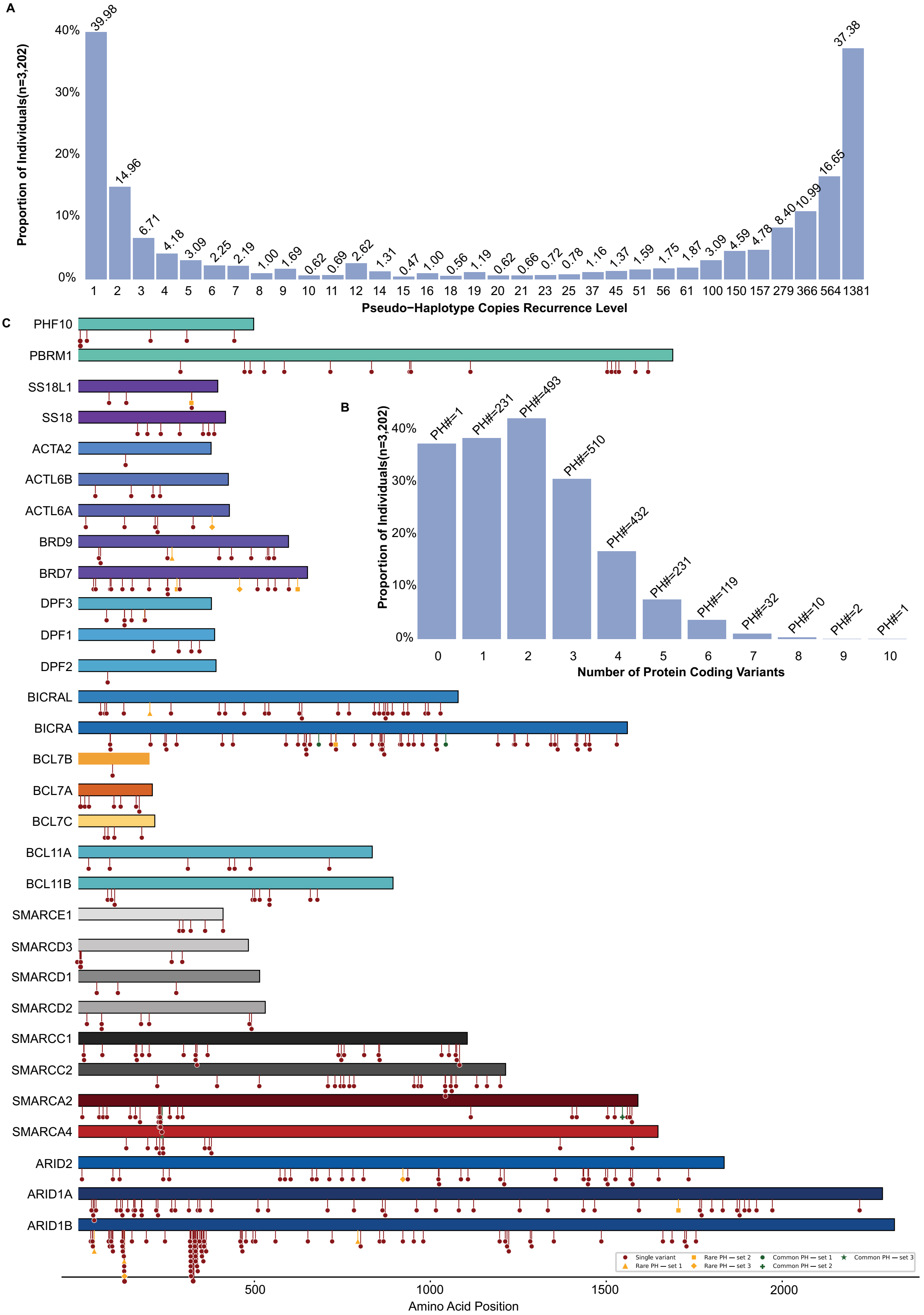


**Figure S1. Individual-level distribution of pseudo-haplotype (PH) carriage.**

**(Fig. 1, continued) (A)** Recurrence-level distribution of individuals (n = 3,202) stratified by protein-coding variant burden. The x-axis shows PH recurrence level (number of identical PH copies), ranging from singletons (recurrence level = 1) to highly recurrent configurations (up to 1,381 copies), and the y-axis represents the proportion of individuals. The distribution is strongly right-skewed, with 39.98% of individuals carrying singleton PHs spanning 1–10 variants. PHs at recurrence levels 2–4 remain common but are enriched for lower variant burdens (primarily 1–3 variants), whereas highly recurrent PHs are shared across larger fractions of individuals and are predominantly low-burden configurations.

**(B; inset)** Distribution of individuals (n = 3,202) by protein-coding variant count. PHs carrying 0–3 variants dominate individual-level carriage (0: 37.38%; 1: 38.44%; 2: 42.22%; 3: 30.61%). Carrier frequency declines sharply with increasing variant burden, with PHs carrying ≥4 variants observed in progressively fewer individuals, indicating constraint against high variant burden at the individual level. Labels indicate the number of distinct pseudo-haplotypes (PH#) within each variant-count category.

**(C)** Distribution of protein-coding variants across amino acid positions in BAF-complex genes. Each horizontal bar represents a gene, scaled to protein length (amino acid position on the x-axis). Red circles indicate individual protein-coding variant sites from the set of 469 curated variants, plotted according to their positions along each protein sequence. Variants are annotated by pseudo-haplotype (PH) classification, including single-variant PHs and multi-variant PHs stratified by frequency-based categories, as indicated by marker shapes and colors. PHs are classified based on the predefined 0.05% PH frequency threshold. Variants are shown across all 31 BAF-complex genes, providing a gene-level overview of their positional distribution along protein sequences.

**Table S1. Protein-coding variants identified through the pseudo-haplotype (PHA) workflow across BAF-complex genes in the 1KGP cohort (n = 469).**

Footnote:

a. This table lists protein-coding and splice-region variants identified across BAF-complex genes in the 1KGP cohort using the pseudo-haplotype (PHA) workflow.

b. Coding variant ID denotes the genomic coordinate-based identifier (chromosome: position: reference: alternate). Coding change (c.) and protein change (p.) are reported using HGVS nomenclature based on Ensembl transcript annotations. For synonymous variants, protein change is denoted as p. (=), indicating no predicted amino acid change. For variants without a defined or predictable protein-level consequence (e.g., splice-site or frameshift variants), protein change is reported as p.(?). In cases where protein annotations are not available in the source data, values are reported as NA. Variant type corresponds to Sequence Ontology (SO) terms describing the functional consequence of each variant. Impact values are derived from CAVA annotations, where 1 indicates high impact and 2 indicates moderate impact.

**Table S2. Integration of GWAS risk alleles with protein-coding variants in BAF-complex genes**

Footnote:

a. This table summarizes genome-wide association study (GWAS) risk alleles mapped to protein-coding variants in BAF-complex genes. GWAS SNPs were obtained from the NHGRI-EBI GWAS Catalog and filtered based on proximity to coding variants within a ±10 kb window to approximate local linkage disequilibrium (LD) structure. Coding variants were derived from 469 curated protein-coding sites identified in the 1000 Genomes Project (1KGP) and annotated using CAVA. Each row represents a GWAS SNP–coding variant pair.

b. Column names are aligned with NHGRI-EBI GWAS Catalog fields, with minor adaptations for consistency. Variant and risk allele corresponds to the SNPS and strongest SNP–Risk Allele fields. P value, P value annotation, OR, Beta, and CI represent GWAS association statistics as reported in the original studies. RAF denotes the risk allele frequency reported in the GWAS study (where available). Mapped genes correspond to MAPPED_GENE. Reported Trait corresponds to DISEASE/TRAIT, and Traits represents the harmonized trait label used in this study. Accession ID, PubMed ID, and Author correspond to GWAS study metadata. Locations indicate the genomic position of the GWAS SNP. Coding variant ID represents protein-coding variants from 1KGP using the format chromosome: position: reference: alternate (e.g., 7:100647509: G:C). Distance to variant indicates the genomic distance (in base pairs) between the GWAS SNP and the corresponding coding variant.

**TableS3.Shared traits across BAF-complex genes based on GWAS risk allele overlap**

Footnote:

a. This table summarizes traits shared across multiple BAF-complex genes based on genome-wide association study (GWAS) risk allele overlap within a ±10 kb window of protein-coding variants, consistent with the filtering criteria described in Table S1. Each row represents a trait for which GWAS SNPs are mapped to two or more BAF-complex genes.

b. Traits denotes the harmonized trait label used in this study. Genes# indicates the number of BAF-complex genes associated with the trait. Variant and risk alleles# represent the number of unique GWAS risk alleles contributing to the trait. Genes lists the BAF-complex genes linked to the trait, and Variants and risk alleles lists the corresponding GWAS SNPs and associated risk alleles (formatted as rsID–allele pairs).

c. Only traits shared by at least two genes are included, representing pleiotropic associations within the BAF complex. Traits shown in † correspond to the representative subset displayed in **Figure 2A**.

**Table S4. Pseudo-haplotype (PH) configurations across BAF-complex genes in the 1KGP cohort**

Footnote:

a. This table lists all 2,062 pseudo-haplotype (PH) configurations constructed from protein-coding variants across 31 BAF-complex genes in the 1000 Genomes Project (1KGP) cohort (n = 6,404 PH copies).

b. Each PH represents a unique combination of co-occurring variants. PH ID denotes the pseudo-haplotype identifier. Pseudo-haplotype identifiers (PH#) are ordered from lowest to highest copy frequency, with higher PH# values indicating more frequent pseudo-haplotypes. Haplotype string (binary) encodes the presence (1) or absence (0) of each variant across a fixed, ordered variant set (see **Table S1**). Constituent variants (HGVS) list all variants within each PH using gene-level HGVS notation (c. and p.). Variant count per PH indicates the number of protein-coding variants per PH (multi-variant burden). PH copy count denotes the number of times each PH is observed in the dataset, and Proportion of PH copies (%) represents the corresponding proportion relative to all PH copies. Carrier samples list individuals harboring each PH; suffixes "_1" and "_2" denote the two chromosomal copies (haplotypes) of each individual.

c.PH IDs marked with † indicate recurrent pseudo-haplotypes observed at or above the 0.05% frequency threshold (≥4 of 6,404 PH copies; n = 122 PHs), corresponding to the subset visualized in **Figure 3A**.

d. The fully reference pseudo-haplotype (0 variants) is represented by PH1 and contains no protein-coding variants.

**Table S5. Multi-variant enrichment analysis of recurrent pseudo-haplotypes across super-populations and the aggregated cohort**

Footnote:

a. This table summarizes the statistical enrichment of multi-variant co-occurrence across pseudo-haplotypes (PHs) identified at the predefined frequency threshold (≥0.05%; ≥4 of 6,404 PH copies) in the 1000 Genomes Project (1KGP) cohort. Analyses were restricted to PHs containing ≥2 protein-coding variants.

b. Column definitions: ID denotes the pseudo-haplotype identifier. Each PH represents a unique combination of co-occurring variants. Pseudo-haplotype identifiers (PH#) are ordered from lowest to highest copy frequency, with higher PH# values indicating more frequent pseudo-haplotypes.

Constituent variants (HGVS) list all protein-coding variants defining each PH using gene-level HGVS nomenclature, including coding (c.) and protein (p.) annotations.

Variant count per PH indicates the number of protein-coding variants comprising each pseudo-haplotype (i.e., multi-variant burden).

Total PH copies (N) denote the total number of pseudo-haplotype copies analyzed within each population (or aggregated cohort).

PH frequency represents the proportion of PH copies carrying the specified variant combination within the given population (Observed / N).

Observed co-occurrences (O) indicates the number of PH copies carrying all variants in the specified combination.

Expected co-occurrences (E) represents the expected number of PH copies carrying the same variant combination under the assumption of independence, calculated from marginal allele frequencies across all PH copies.

Expected probability (P_null) denotes the expected joint probability of observing the variant combination under the independence model (product of marginal allele frequencies).

Pseudo-count applied indicates whether a pseudo-count (0.5) was applied to observed and expected counts to ensure numerical stability in enrichment calculations.

$\text{Log}_2$ enrichment quantifies deviation from independence, where positive values indicate co-occurrence (enrichment) and negative values indicate mutual exclusivity (depletion).

Lower 95% CI and Upper 95% CI represent the confidence interval bounds for the $\log_2$ enrichment estimate.

P-value denotes the statistical significance of deviation from independence, calculated using a two-sided exact binomial test.

FDR-adjusted P-value represents the P-value corrected for multiple testing using the Benjamini–Hochberg procedure.

Association type classifies each variant combination as co-occurrence ($\log_2$ enrichment $> 0$ and FDR $< 0.05$), mutual exclusivity ($\log_2$ enrichment $< 0$ and FDR $< 0.05$), or not significant (FDR $\geq 0.05$).

Population indicates the super-population in which the analysis was performed—African (AFR), American (AMR), East Asian (EAS), European (EUR), or South Asian (SAS)—or the aggregated cohort.

c. PH IDs marked with † indicate pseudo-haplotypes visualized in Figure.4A–B for the aggregated population. PH IDs marked with * correspond to pseudo-haplotypes shown in Figure.5 across the five super-populations and the aggregated population.
